# Modulation of the combinatorial code of odorant receptor response patterns in odorant mixtures

**DOI:** 10.1101/865386

**Authors:** Claire A. de March, William B. Titlow, Tomoko Sengoku, Patrick Breheny, Hiroaki Matsunami, Timothy S. McClintock

**Affiliations:** Department of Molecular Genetics and Microbiology, Duke University School of Medicine, Durham, NC 27710, USA; Department of Physiology, University of Kentucky, Lexington, Kentucky 40536-0298, USA; Department of Biostatistics, University of Iowa, Iowa City, IA 52242, USA; Department of Neurobiology, Duke University School of Medicine, Durham, NC 27710, USA; Institute of Global Innovation Research, Tokyo University of Agriculture and Technology, Koganei, Tokyo 184- 8588, Japan; Duke Institute for Brain Sciences, Duke University, Durham, NC 27710, USA

**Keywords:** olfactory coding, olfactory sensory neurons, G protein-coupled receptors, olfaction

## Abstract

The perception of odors relies on combinatorial codes consisting of odorant receptor (OR) response patterns to encode odor identity. The modulation of these patterns by odorant interactions at ORs potentially explains several olfactory phenomena: mixture suppression, unpredictable sensory outcomes, and the perception of odorant mixtures as unique objects. We determined OR response patterns to 4 odorants and 3 binary mixtures *in vivo* in mice, identifying 30 responsive ORs. These patterns typically had a few strongly responsive ORs and a greater number of weakly responsive ORs. The ORs responsive to an odorant were often unrelated sequences distributed across several OR subfamilies. Mixture responses predicted pharmacological interactions between odorants, which were tested *in vitro* by heterologous expression of ORs in cultured cells. These tests provided independent evidence confirming odorant agonists for 13 ORs and identified both suppressive and additive effects of mixing odorants. This included 11 instances of antagonism of ORs by an odorant, 1 instance of additive responses to a binary mixture, 1 instance of suppression of a strong agonist by a weak agonist, and the discovery of an inverse agonist for an OR. These findings demonstrate that interactions between odorants at ORs are common.

## Introduction

In terrestrial environments, odors are comprised of volatile chemicals called odorants. The ability to detect odorants evolved via diversification and specialization of plasma membrane receptors that are expressed primarily in the olfactory sensory neurons (OSNs) of the olfactory epithelium. For mammals, the detectors are G-protein coupled receptors (GPCRs), primarily those of the odorant receptor (OR) family, but the much smaller family of trace amine-associated receptors (TAARs) contributes by helping to detect odorants containing amines (Liberles and Buck, 2006; Johnson et al., 2012). In most mammalian species, ORs are the largest family of genes in the genome, with ∼1,100 intact OR genes in mice, ∼400 in humans and nearly 2,000 in elephants, for example (Niimura et al., 2014). When an odorant agonist binds an OR, the OR favors an active conformation that stimulates a heterotrimeric G-protein containing Gα_olf_. This in turn activates adenylyl cyclase to produce the second messenger molecule cyclic adenosine 3’-monophosphate (cAMP). It causes an electrical response via the successive activation of a cation-permeable cyclic nucleotide-gated ion channel and a calcium-activated chloride channel (Kleene, 2008). Each OSN expresses just one allele of one OR gene (Chess et al., 1994; Malnic et al., 1999), so the pharmacology of one OR determines the response properties of each OSN. Furthermore, expression of a single allele of one OR gene allows the OSNs innervating each glomerulus in the olfactory bulb to be restricted to those expressing the same OR. Consequently, patterns of OR response are faithfully transmitted to the brain in the form of spatiotemporal patterns of glomerular activity (Treloar et al., 2002). This organization offers an elegant explanation for how the olfactory periphery provides the information needed for odor discrimination and the perception of odor qualities. The critical nature of OR response patterns is evident. When the OR or TAAR most sensitive to an odorant is genetically deleted, or when a large majority of OSNs are forced to express the same OR, the ability to detect cognate odorants and discriminate them from other odorants is diminished (Fleischmann et al., 2008; Sato-Akuhara et al., 2016; Dewan et al., 2018; Horio et al., 2019). At present, relatively little is known about the nature of OR response patterns *in vivo* (McClintock et al., 2014; Jiang et al., 2015; von der Weid et al., 2015; Dewan et al., 2018).

Natural odors are usually mixtures of odorants, so the potential for interactions at ORs nearly always exists when an odor is encountered. *In vitro* heterologous expression studies of ORs in cultured cells and *ex vivo* experiments on dissociated OSNs have demonstrated that odorants are often capable of acting as antagonists or inverse agonists at some ORs (Oka et al., 2004; Sanz et al., 2005; Reisert, 2010; Reddy et al., 2018) even while they act as agonists at other ORs (Araneda et al., 2000; Spehr et al., 2003; Oka et al., 2004; Sanz et al., 2008). These results are potential explanations for long-standing evidence of perceptual and physiological interactions between odorants in animals ranging from arthropods to mammals (Thomas-Danguin et al., 2014). Even odor pleasantness, a perceptual dimension more reliably linked to odorant structure than most descriptors of odor sensations (Keller and Vosshall, 2016), is not consistently predictable from the pleasantness of the components of odor mixtures (Lindqvist et al., 2012). The components of odorant mixtures are sometimes difficult to identify even in binary mixtures, though training improves performance on such tasks (Rokni et al., 2014), and humans have increasing difficulty identifying the components of mixtures as mixture complexity increases (Laing and Francis, 1989; Livermore and Laing, 1996; Poupon et al., 2018). Further work on this problem has confirmed that olfactory signal processing is capable of identifying individual odorant components in certain situations, but often odor mixtures are perceived instead as unique odor objects (Thomas-Danguin et al., 2014). This synthesis of odor objects is believed to derive not only from central olfactory processing but also from peripheral events such as odorant interactions at ORs (Cromarty and Derby, 1998; Rospars et al., 2008; Munch et al., 2013). These odorant interactions could bias the olfactory system towards perceiving odor mixtures as distinct odor objects by obscuring the combinatorial codes of the component odorants due to suppressive interactions that eliminate parts of their OR response patterns and introducing novel responses from ORs when additive or synergistic interactions occur (Figure 1A).

**Figure 1.**
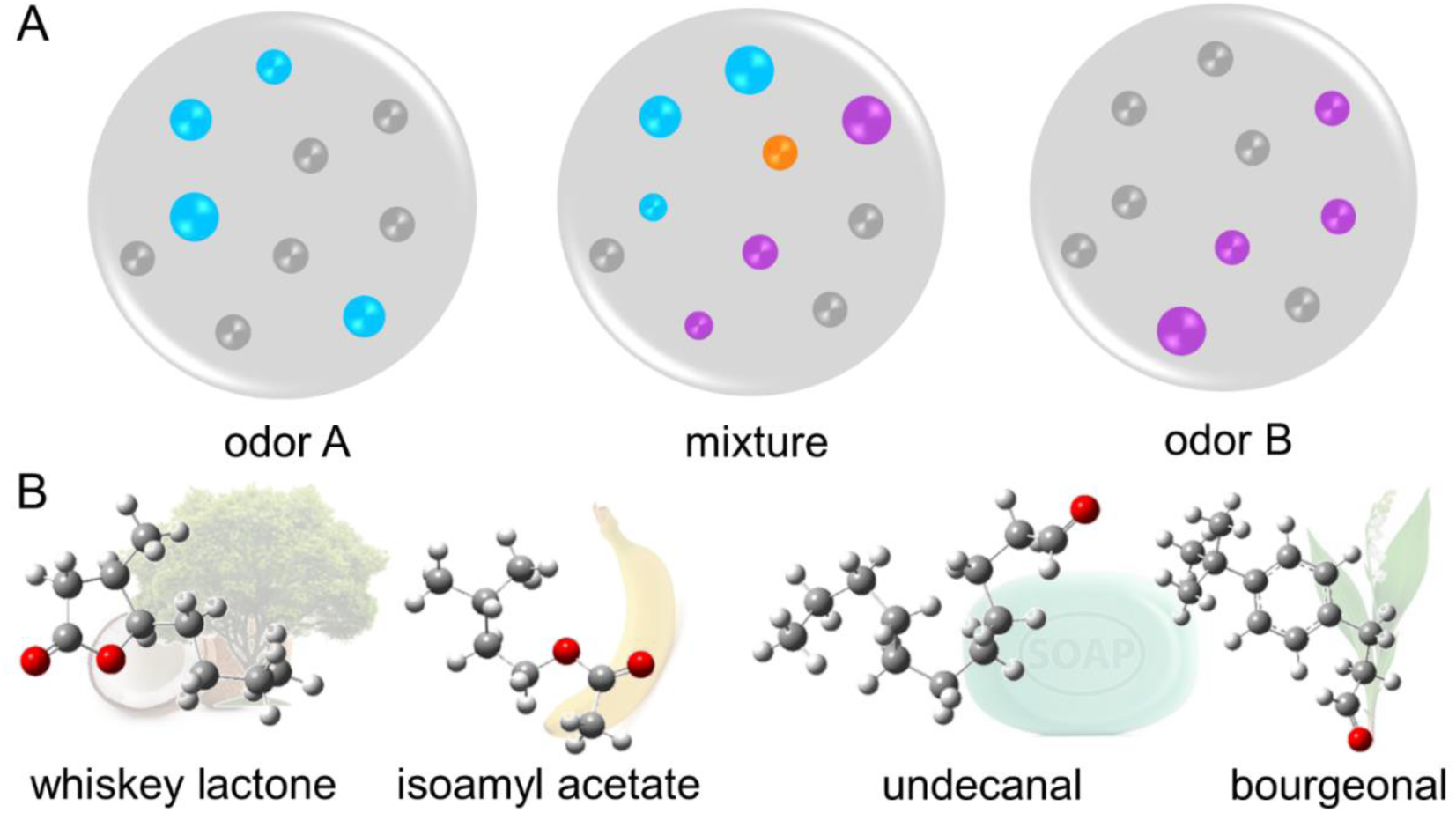
Odor coding theory and odorant compounds investigated. (**A**) Schematic representation of the modulation of the combinatorial code of activated ORs. ORs are represented by dots (gray signifies no response) and their size indicates response strength. Odor A activates a pattern of ORs (blue) and odor B activates a different pattern (violet). The mixture of odor A and B shows a response pattern that is not the sum of the responses to odors A and B, with some ORs continuing to respond (blue and violet, respectively), some that are suppressed (gray), and some that respond only to the mixture due to an additive or synergistic effect (orange). (**B**) Tridimensional structures of the four compounds used in this study with their associated odor percept. Carbon atoms are represented in gray, hydrogen in white and oxygen in red.

We used our ability to measure OR function *in vivo* (McClintock et al., 2014) to identify OR responses to binary odor mixtures, including two mixtures whose component odorants are known to interact at perceptual or physiological levels (Figure 1B). One mixture with a known interaction, inspired by wine flavor complexity, is composed of isoamyl acetate and whiskey lactone, which have fruity and woody odors, respectively. In mixtures containing high concentrations of whiskey lactone, the perception of the fruity odor of isoamyl acetate is suppressed (Chaput et al., 2012). Another mixture with a known interaction, initially identified in the context of the chemotaxis of human sperm, is bourgeonal and undecanal, which have floral and soapy odors, respectively (Spehr et al., 2003). Later, undecanal was shown to decrease the perception of bourgeonal in mixtures of these two odorants (Brodin et al., 2009). Furthermore, mice are unusually sensitive to bourgeonal in behavioral assays, making this odorant worthy of detailed investigation (Larsson and Laska, 2011). Once OR responses to these odorants and their mixtures were identified *in vivo*, we used our *in vitro* assay (Zhang et al., 2017) to investigate the pharmacology of individual ORs. We found that OR responses to an odorant are often different when a second odorant is present. These data demonstrate that pharmacological interactions between odorants at ORs can have substantial effects on OR response patterns, confirming modulation of the combinatorial code during the perception of a mixture of odorants.

## Results

### Odorant receptors responsive to isoamyl acetate

To identify ORs responsive to isoamyl acetate odorants we used an *in vivo* assay that simultaneously measures all ORs and TAARs (McClintock et al., 2014). This assay uses odor-stimulated expression of green fluorescent protein (GFP) from the activity-dependent *S100A5* gene locus in S100a5-tauGFP mice. This sequence of events was demonstrated in double mutant mice where the absence of Cnga2 prevents olfactory transduction from generating a receptor potential and consequently prevents expression of GFP from the *S100a5* locus (McClintock et al., 2014). Therefore, transcription from this locus and expression of the GFP reporter rely on electrical activity resulting from OR activity. In S100a5-tauGFP mice stimulated with headspace air above solutions of either diluted odorants or diluent alone fluorescence-activated cell sorting (FACS) is used to capture samples of GFP^+^ and GFP^-^ cells dissociated from olfactory epithelia. High-throughput expression profiling techniques allow simultaneous screening of all ORs and TAARs to identify OR or TAAR mRNAs enriched in GFP^+^ FACS samples relative to GFP^-^ FACS samples from mice stimulated with odorant, but not enriched in GFP^+^ FACS samples from control mice. These specifically enriched mRNAs must encode ORs or TAARs responsive to the tested odorant because OSNs express only one OR or TAAR per OSN.

We first tested the headspace air above a 5% solution of isoamyl acetate. Isoamyl acetate evoked responses from 14 ORs (Figure 2A), including strong responses of more than 3-fold from 6 of these ORs (Table S1). To confirm that these *in vivo* data identify ORs responsive to isoamyl acetate we used our *in vitro* method of heterologous expression in cultured cells (Zhang et al., 2017) to test 13 of these ORs. This Glosensor *in vitro* assay utilizes Gnal to allow ORs expressed in cultured cells to activate cAMP signaling, the same pathway activated *in vivo*. The conservative nature of the *in vivo* assay, which delivers odorants intermittently at concentrations less than saturation, gives reason to expect false negatives so we also tested 2 ORs that approached significance *in vivo*. We first did a simple *in vitro* screen using 1000 µM isoamyl acetate. This screen identified 5 responsive ORs, including 4 that responded *in vivo* (Olfr213, Olfr1411, Olfr183, Olfr1126), and one OR (Olfr167) that gave a nearly significant response *in vivo* (Table S1). Next, we measured isoamyl acetate dose-response relationships *in vitro* for these 5 ORs. The *in vitro* data from Olfr1126 are equivocal (Figure 2P), indicating that it is at best weakly responsive to isoamyl acetate, but these experiments confirm strong responses to isoamyl acetate from Olfr183, Olfr213, Olfr1411, and Olfr167 (Figure 2G, J, M, and S). The agreement between *in vivo* and *in vitro* data demonstrates unequivocally that isoamyl acetate is an agonist for Olfr183, Olfr213, Olfr1411, and Olfr167.

**Figure 2.**
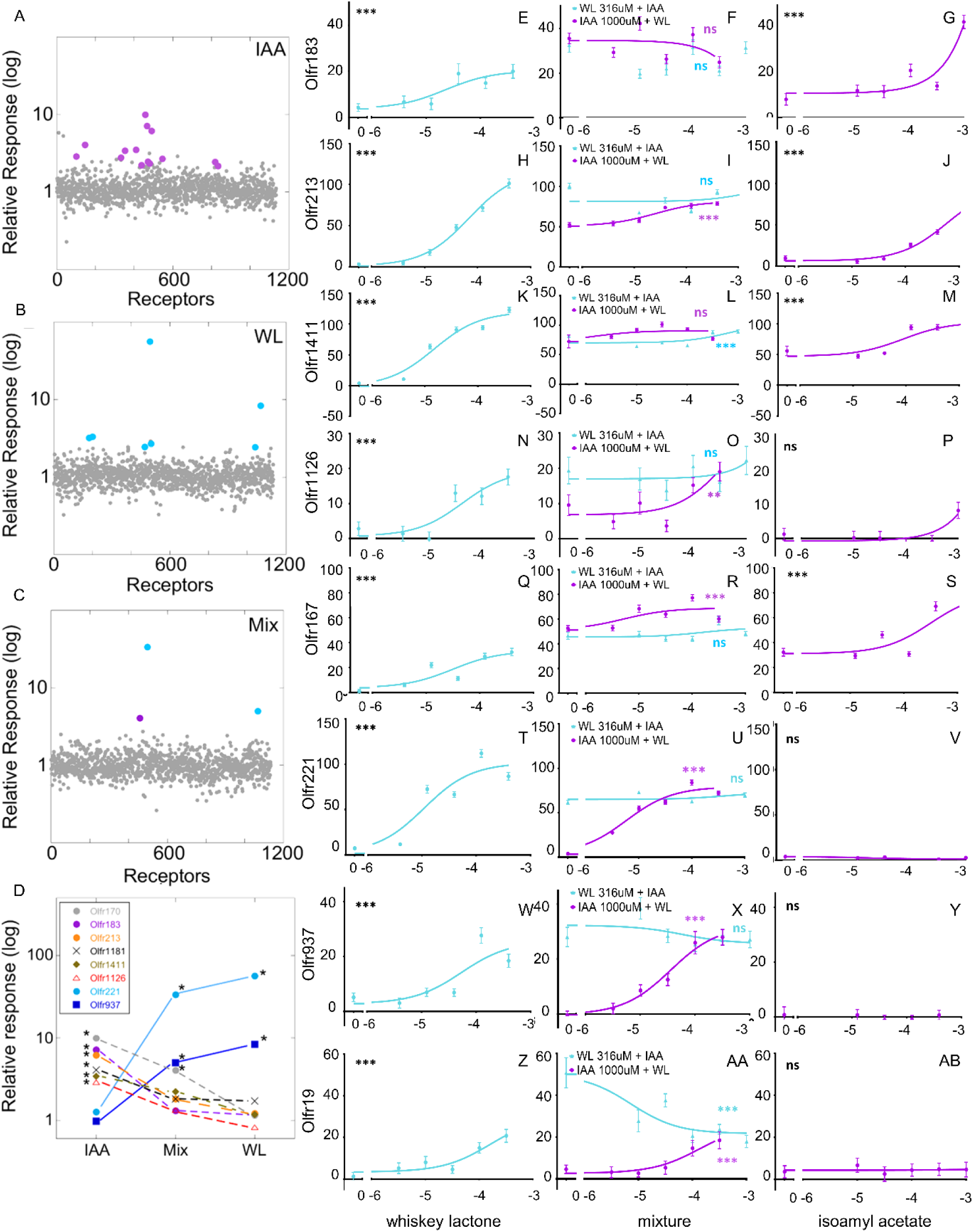
Odorant receptors responsive to isoamyl acetate (IAA) and whiskey lactone (WL). Of the more than 1,100 ORs and trace-amine associated receptors (TAARs) expressed in the mouse olfactory epithelium (shown in alphabetical order on the x-axis in **A-C**), 14 ORs responded to 5% IAA (violet) *in vivo* (**A**) and 7 ORs responded to 50% WL (blue) *in vivo* (**B**). Only the 2 ORs highly responsive to WL and the 1 OR most responsive to IAA show significant responses to the mixture of IAA and WL (**C**). Comparing the *in vivo* response magnitudes (**D**) reveals consistently negative interactions of IAA and whiskey lactone WL at ORs. The 6 ORs most responsive to IAA show reduced responses when WL is present (Mix), and only Olfr170 continues to show a significant response. The other 5 ORs fail to respond to the mixture of IAA and WL (Mix). In contrast, the 2 ORs most responsive to WL continue to respond strongly to the mixture. (*), significant response (FDR ≤ 10%). *In vitro* measures of responses to whiskey lactone (blue) in panels **E, H, K, N, Q, T, W, and Z** and isoamyl acetate (violet) in panels **G, J, M, P, S, V, Y** and **AB**. Binary mixtures of increasing doses of one odorant mixed with a consistent dose of the other odorant are shown in panels **F, I, L, O, R, U, X**, and **AA**. In the *in vitro* assay plots the x-axes represent the log of the odorant concentration (M) and the y-axes show the normalized Glosensor luminescence response to cAMP. ANOVA trend analysis *p<0.05; **p<0.01; ***p<0.001.

Agreement between *in vivo* and *in vitro* methods provides certainty about the nature of odorant ligand effects on specific ORs. Lack of agreement does not lead to similarly firm conclusions, however, because lack of agreement can arise from limitations in both assays. For example, the *in vivo* assay does not deliver odorants at concentrations as high as is possible in the *in vitro* assay, so weakly sensitive ORs may be detected poorly *in vivo*. For the *in vitro* assay a well-known technical limitation when expressing ORs in heterologous cells is poor plasma membrane trafficking of some ORs (McClintock et al., 1997; Gimelbrant et al., 2001; McClintock and Sammeta, 2003; Saito et al., 2009) so we measured the level of cell surface expression of several ORs during heterologous expression. We compared these levels against those of an OR that traffics well, Olfr539, and an OR that traffics poorly, Olfr541 (Ikegami et al., 2019). Olfr213, Olfr1411, and Olfr1126 were detected in the plasma membrane of HEK293 cells at levels greater than Olfr541 and all responded to isoamyl acetate (Figure S1). Olfr170, Olfr1181, Olfr183, and Olfr1480 showed lower levels of plasma membrane expression than Olfr541. While this might explain why Olfr170, Olfr1181, and Olfr1480 responded *in vivo* but failed to respond *in vitro*, the ability of Olfr183 to respond to isoamyl acetate *in vitro* argues that low expression does not necessarily prevent functionality in this assay system, as previously noted in the case of OR7D4 (de March et al., 2018).

### Odorant receptors responsive to whiskey lactone

Whiskey lactone evoked responses from 7 ORs *in vivo*, including strong responses from 2 of them (Figure 2B, Table S1). These 2 most responsive ORs, Olfr221 and Olfr937, also responded to whiskey lactone *in vitro*, as did Olfr19 (Figure 2T, W, and Z). Measuring the level of cell surface expression revealed that Olfr221 traffics better than Olfr541, an OR known to traffic poorly, while Olfr19 and Olfr937 showed lower levels of plasma membrane expression than Olfr541 (Figure S1). That Olfr19 and Olfr937 were functional provides further evidence that the level of plasma membrane expression is an imperfect predictor of functionality in our *in vitro* system (de March et al., 2018).

### Cross-sensitivity of odorant receptors to isoamyl acetate and whiskey lactone

Isoamyl acetate and whiskey lactone interact perceptually in humans and physiologically at rat OSNs (Chaput et al., 2012). A likely explanation for these data is that the odorants interact at certain ORs, altering the pattern of OR responses in ways that would cause a change in OSN input to the brain and odor perception. To investigate the initial step in this hypothesis we tested the headspace air above a mixture of isoamyl acetate (5%) and whiskey lactone (50%). *In vivo*, this mixture evoked responses from only a subset of the ORs responsive to these two odorants (Figures 2C and 2D, Table S1). The two ORs that responded most strongly to whiskey lactone, Olfr221 and Olfr937, continued to respond strongly to the binary mixture with at most a small decrease in response, but the five weakly responsive ORs no longer showed a significant response (Figures 2C and 2D, Table S1). The ORs responsive to isoamyl acetate were even more strongly affected by the mixture (Figures 2C and 2D, Table S1). Of the ORs responsive to isoamyl acetate *in vivo*, only the most responsive OR, Olfr170, continued to give a significant response. These data are consistent with the ability of whiskey lactone to suppress isoamyl acetate responses from 43% of freshly isolated rat OSNs that respond to isoamyl acetate, and to suppress electroolfactogram responses to isoamyl acetate in rats (Chaput et al., 2012).

When we used a mixture of 1000 µM isoamyl acetate and 316 µM whiskey lactone to do an *in vitro* screen of 21 ORs for mixture effects we observed strong responses from the 8 ORs that respond to either isoamyl acetate or whiskey lactone, as described above, and from Olfr1480 (Table S1). Measuring dose-response relationships for isoamyl acetate and whiskey lactone confirmed that Olfr221, Olfr937, and Olfr19 respond to whiskey lactone but not to isoamyl acetate (Figure 2T, V, W, Y, Z, and AB). However, the 5 ORs responsive to isoamyl acetate *in vitro* also responded to whiskey lactone (Figure 2E, B, H, J, K, M, N, P, Q, S). To more thoroughly investigate the sensitivity of these ORs to these two odorants we designed an *in vitro* experiment capable of simultaneously testing for both dose-dependent antagonism and differences in agonist efficacy. When testing a consistent dose of one odorant mixed with a series of increasing concentrations of another odorant, antagonism results in a dose-dependent decrease in response magnitude while differences in efficacy result in a dose-dependent increase in response magnitude above that of the odorant held at a consistent dose. This experiment further confirmed that whiskey lactone, but not isoamyl acetate, was an agonist for Olfr221, Olfr937, and Olfr19 (Figure 2U, X, AA). Responses of Olfr221 and Olfr937 to whiskey lactone were not sensitive to isoamyl acetate (Figure 2U, X) but isoamyl acetate was an antagonist of responses to whiskey lactone at Olfr19 (Figure 2AA). Isoamyl acetate and whiskey lactone had similar efficacy at Olfr183 (Figure 2F). Isoamyl acetate and whiskey lactone also had similar efficacy at Olfr1411, though isoamyl acetate may be a slightly better agonist (Figure 2L). In contrast, whiskey lactone was a stronger agonist than isoamyl acetate at Olfr213, Olfr1126, and Olfr167 (Figure 2I, O, and R). These data confirmed that most ORs responsive to whiskey lactone were insensitive to isoamyl acetate but that many ORs responsive to isoamyl acetate were sensitive to whiskey lactone. The observation that the whiskey lactone sensitivity of ORs responsive to isoamyl acetate takes the form of agonism *in vitro* rather than antagonism is identical to heterologous expression data from two human ORs, which responded both to isoamyl acetate and to whiskey lactone (Chaput et al., 2012).

### Odorant receptors responsive to undecanal or bourgeonal

Undecanal has been reported to be an antagonist of a human OR that responds to bourgeonal in an *in vitro* assay and to suppress the perception of bourgeonal by humans when undecanal is at higher concentrations than bourgeonal (Spehr et al., 2003; Brodin et al., 2009). Mice are unusually sensitive to bourgeonal, responding to lower concentrations than other odorants in behavioral assays (Larsson and Laska, 2011). These findings suggest that like isoamyl acetate and whiskey lactone the mixture of these two odorants might show interesting patterns of OR responses, including interaction effects at specific ORs.

When we tested undecanal (headspace above a 5% solution) *in vivo* in mice, we detected significant responses from 2 ORs: Olfr774 and Olfr1419 (Figure 3A, Table S2). *In vitro* testing confirmed that Olfr774 responded to undecanal but not to bourgeonal (Figure 3E, G). When tested *in vivo* in mice, bourgeonal (headspace above a 2% solution) evoked responses from 7 ORs, including an especially strong response from Olfr16 – one of the largest responses we have yet observed (Figure 3B, Table S2). We also detected a strong response to bourgeonal from Olfr1099 and weaker responses from Olfr1040, Olfr1151, Olfr198, Olfr1049, and Olfr738. *In vitro* screening of these ORs with 100 µM bourgeonal detected responses from Olfr16 and Olfr1099 (Table S2). Measurements of dose-response relationships confirmed that bourgeonal was a strong agonist for Olfr16 and that undecanal had no agonist activity at this OR (Figure 3H-J).

**Figure 3.**
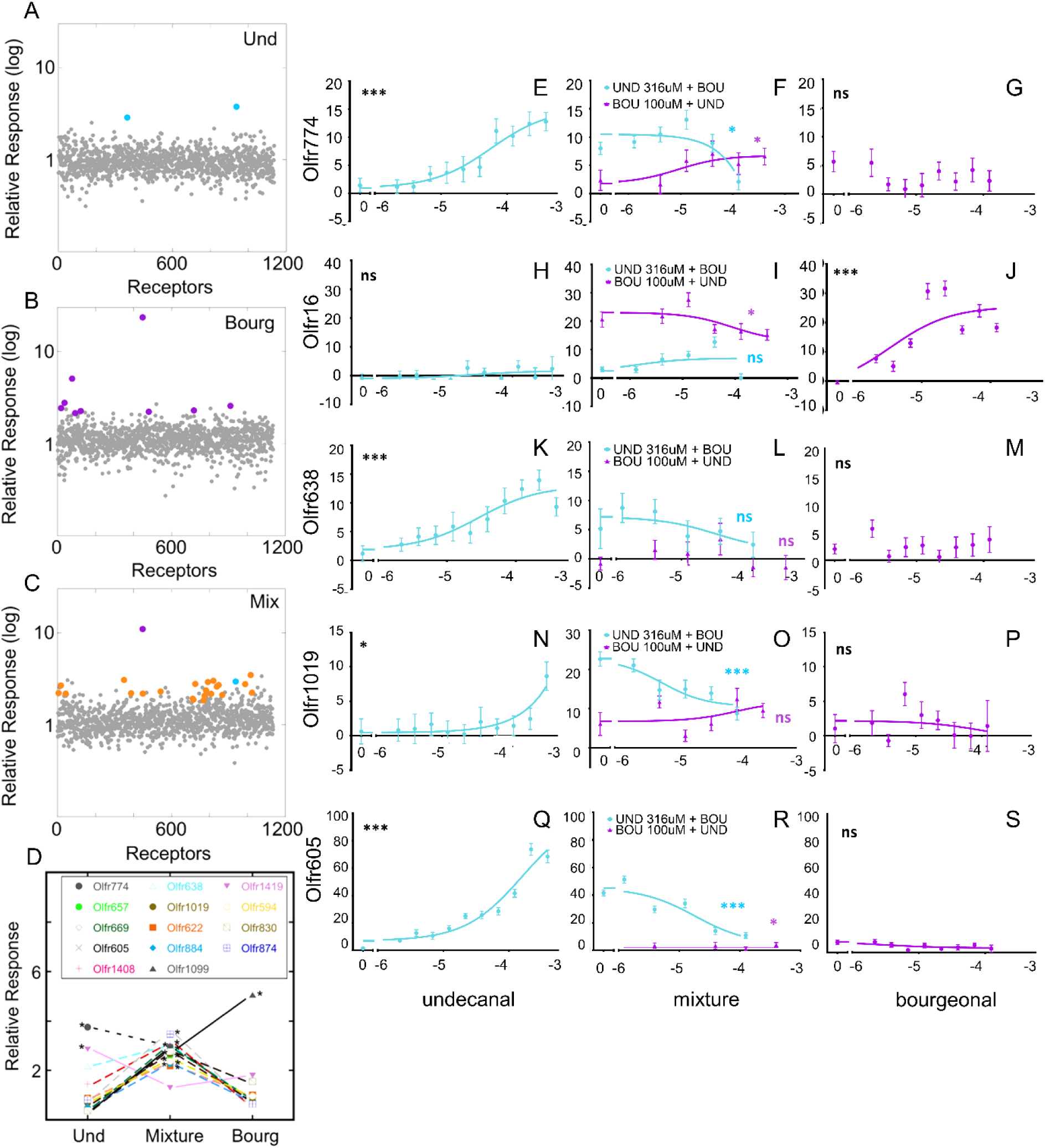
OR response patterns to bourgeonal and undecanal. 2 ORs respond to 5% undecanal *in vivo* (**A**, blue) and 9 ORs respond to 2% bourgeonal (**B**, violet) and 24 ORs respond to the mixture of bourgeonal and undecanal (**C**), including the 2 ORs most responsive to bourgeonal and undecanal, respectively, and 22 ORs that respond specifically to the mixture (orange). X-axis, ORs and TAARs in alphabetical order. Two patterns of changes in OR response magnitudes occur *in vivo* when bourgeonal and undecanal are mixed (**D**). The most responsive ORs show decreased responses to the mixture. (For clarity, the massive responses of Olfr16 to bourgeonal and to the mixture are omitted from the plot.) Mixture-specific ORs are more responsive to the mixture than to either odorant alone. (*), significant response (FDR ≤ 10%). *In vitro* measures of responses to undecanal (blue) in panels **E, H, K, N, and Q**, and bourgeonal (violet) in panels **G, J, M, P, and S**. Binary mixtures of increasing doses of one odorant mixed with a consistent dose of the other odorant are shown in panels **F, I, L, O, and R**. In the *in vitro* assay plots the x-axes represent the log of the odorant concentration (M) and the y-axes show the normalized Glosensor luminescence response to cAMP. ANOVA trend analysis *p<0.05; ***p<0.001.

### Interactions between bourgeonal and undecanal at odorant receptors

When we did *in vivo* tests of a mixture of bourgeonal and undecanal the ORs strongly and specifically responsive to bourgeonal or undecanal showed smaller but significant, or nearly significant, responses (Figure 3C-D). The pattern of reduced response magnitudes of Olfr774 and Olfr16 to the mixture predict antagonist effects of undecanal and bourgeonal, respectively, at these ORs. *In vitro* tests confirmed that bourgeonal was a weak antagonist of Olfr774 responses to undecanal (Figure 3F) and that undecanal was a weak antagonist of Olfr16 responses to bourgeonal (Figure 3I).

The *in vivo* response to the mixture also included weak responses from 22 additional ORs that failed to reach significance in tests of bourgeonal or undecanal alone (Figure 3C). These ORs had greater magnitudes of response to the mixture than to bourgeonal or undecanal alone, raising the possibility of additive or synergistic responses (Figure 3D, Table S2). To test this idea we performed *in vitro* assays. We evaluated the cell surface expression of 8 of these ORs and screened 19 of them for function using 316 µM undecanal and 100 µM bourgeonal. Olfr1019, Olfr16, and Olfr638 were expressed at higher levels in the plasma membrane than the low expression control, Olfr541, while Olfr1099, Olfr577, Olfr1420, Olfr605 and Olfr774 showed lower expression than Olfr541 (Figure S1). The *in vitro* screen revealed responses to one or both odorants from Olfr638, Olfr605, Olfr1019, and Olfr1420 (Table S2). These ORs were selected for more detailed study.

Measuring dose-response relationships confirmed that Olfr638, Olfr1019, and Olfr605 responded to undecanal but not to bourgeonal (Figure 3K, M, N, P, Q, and S). Using the odorant mixture test for dose-dependent antagonism and differences in agonist efficacy, we evaluated whether bourgeonal and undecanal interact at these 3 ORs. Undecanal responses of Olfr638 were insensitive to bourgeonal but responses of Olfr1019 and Olfr605 to undecanal were antagonized by bourgeonal (Figure 3L, O, and R). The selective response of these ORs to the mixture of undecanal and bourgeonal *in vivo* was not due to some type of additive response, but rather appears to have been variation in the detection of weak responses.

In contrast to the agonist specificity of the other ORs sensitive to the mixture of bourgeonal and undecanal, Olfr1420 responded to both bourgeonal and undecanal *in vitro*, with undecanal evoking stronger responses (Figure 4A-C, Tables S2 and S3). Unlike the *in vitro* data showing additive responses of several ORs that responded well to both whiskey lactone and isoamyl acetate, the large difference in efficacy between bourgeonal and undecanal at Olfr1420 required a more sensitive approach to adequately test for an additive response *in vitro*. We used an orthogonal array of mixtures of concentrations of bourgeonal and undecanal to confirm that Olfr1420 shows additive responses to mixtures of bourgeonal and undecanal (Figure 4D; Tables S2 and S3). The ability of Olfr1420 to respond to both bourgeonal and undecanal explained why this OR would show evidence of an additive response to mixtures of these two odorants *in vivo*. The Olfr1420 data also confirmed a prediction, based on modeling of OSN response patterns, that one mechanism of suppressive interactions between odorants is the ability of partial agonists to suppress responses from stronger agonists (Reddy et al., 2018). We observed a dose-dependent decrease in the response from Olfr1420 when increasing concentrations of the weak agonist bourgeonal were mixed with a consistent high concentration (316µM) of the strong agonist undecanal (Figure 4C).

**Figure 4.**
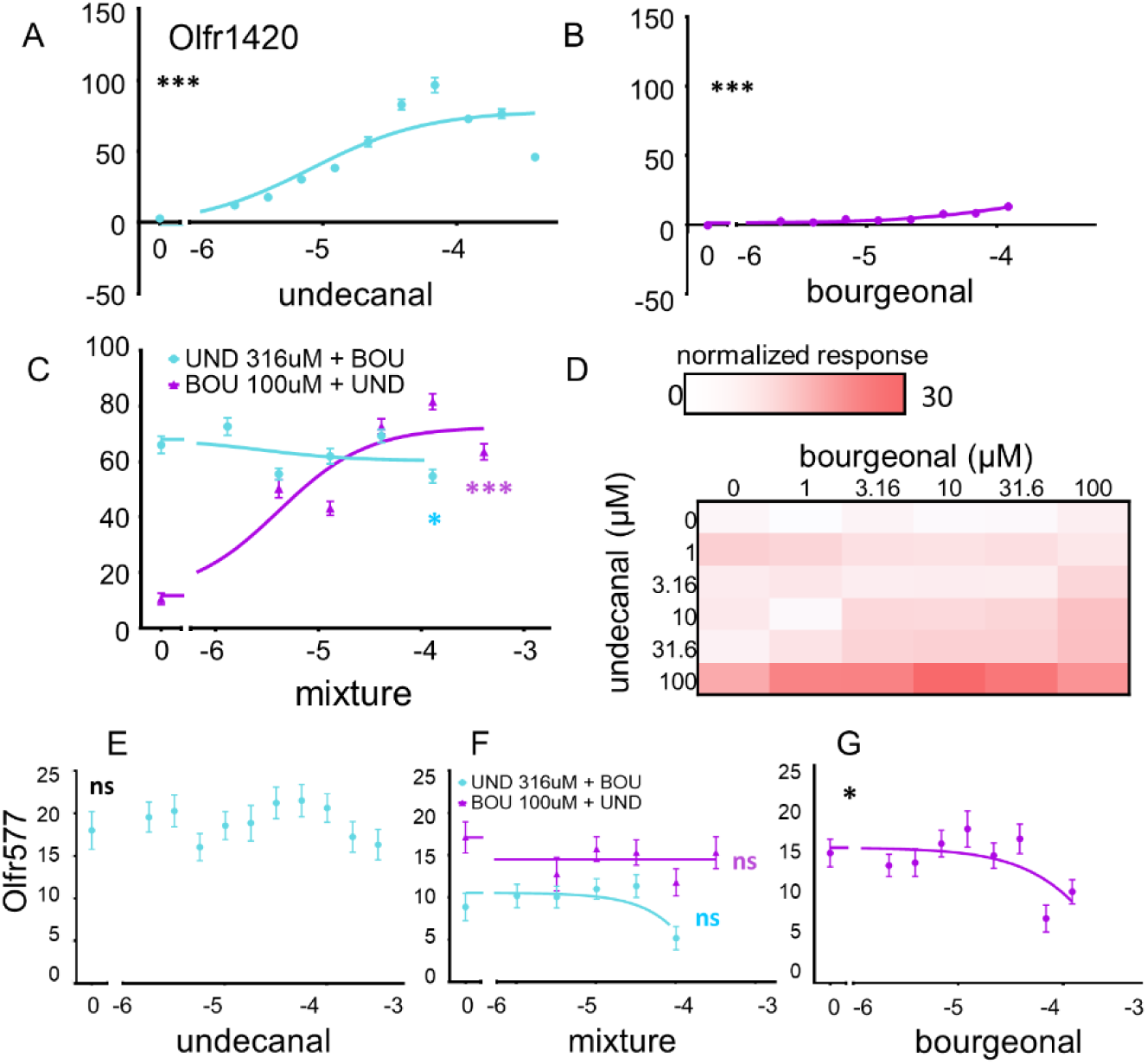
Additive interaction of undecanal and bourgeonal at Olfr1420 and inverse agonism of bourgeonal at Olfr577. Dose-response relationships of Olfr1420 to undecanal (blue) (**A**) and bourgeonal (violet) (**B**) and binary mixtures (**C**). (C) The strong agonist undecanal causes a dose-dependent increase in response on top of the weak response to 316µM bourgeonal (violet), and the weak agonist bourgeonal causes a dose-dependent suppression of the response to 100µM of the strong agonist undecanal (blue). (**D**) Additive effects of undecanal and bourgeonal are also apparent in the matrix of Olfr1420 responses to an orthogonal array of concentrations of binary mixtures of these odorants. The normalized response is depicted as white for no response grading to red for the maximum response (30). See Supplementary Table S3 for response magnitude values. Dose-response relationship of Olfr577 to undecanal (blue) (**E**) and bourgeonal (violet) (**G**) and binary mixtures (**F**). ANOVA trend analysis *p<0.05; **p<0.01; ***p<0.001.

### Bourgeonal is an inverse agonist for Olfr577

ORs are thought to have varying levels of constitutive activity (Reisert, 2010), and when this activity is high it provides an opportunity to discriminate odorants acting as antagonists from those acting as inverse agonists, as has been done previously in one instance (Reisert, 2010). We discovered an instance of inverse agonism involving Olfr577, which showed substantial odorant-independent constitutive activity, also called basal activity, in our *in vitro* assay (Figure 4E-G). Olfr577 was insensitive to undecanal (Figure 4E) but bourgeonal significantly decreased the basal signal from this OR (p=0.017) (Figure 4G).

### Interactions between whiskey lactone and undecanal at odorant receptors

Interactions at ORs between odorants known to interact perceptually proved to be common, so to extend the analysis to odorants not known previously to interact perceptually, we tested a binary mixture of whiskey lactone and undecanal. These two odorants are not related structurally and their woody and soapy percepts, respectively, do not appear to be related (Figure 1B). When we tested the headspace above of a mixture of 5% undecanal and 50% whiskey lactone we observed that most ORs responsive to these two odorants gave reduced responses to the mixture *in vivo* (Figure 5A - D). Only Olfr711, which showed evidence suggestive of a response to whiskey lactone alone (FDR = 0.20), showed a stronger response to the mixture.

**Figure 5.**
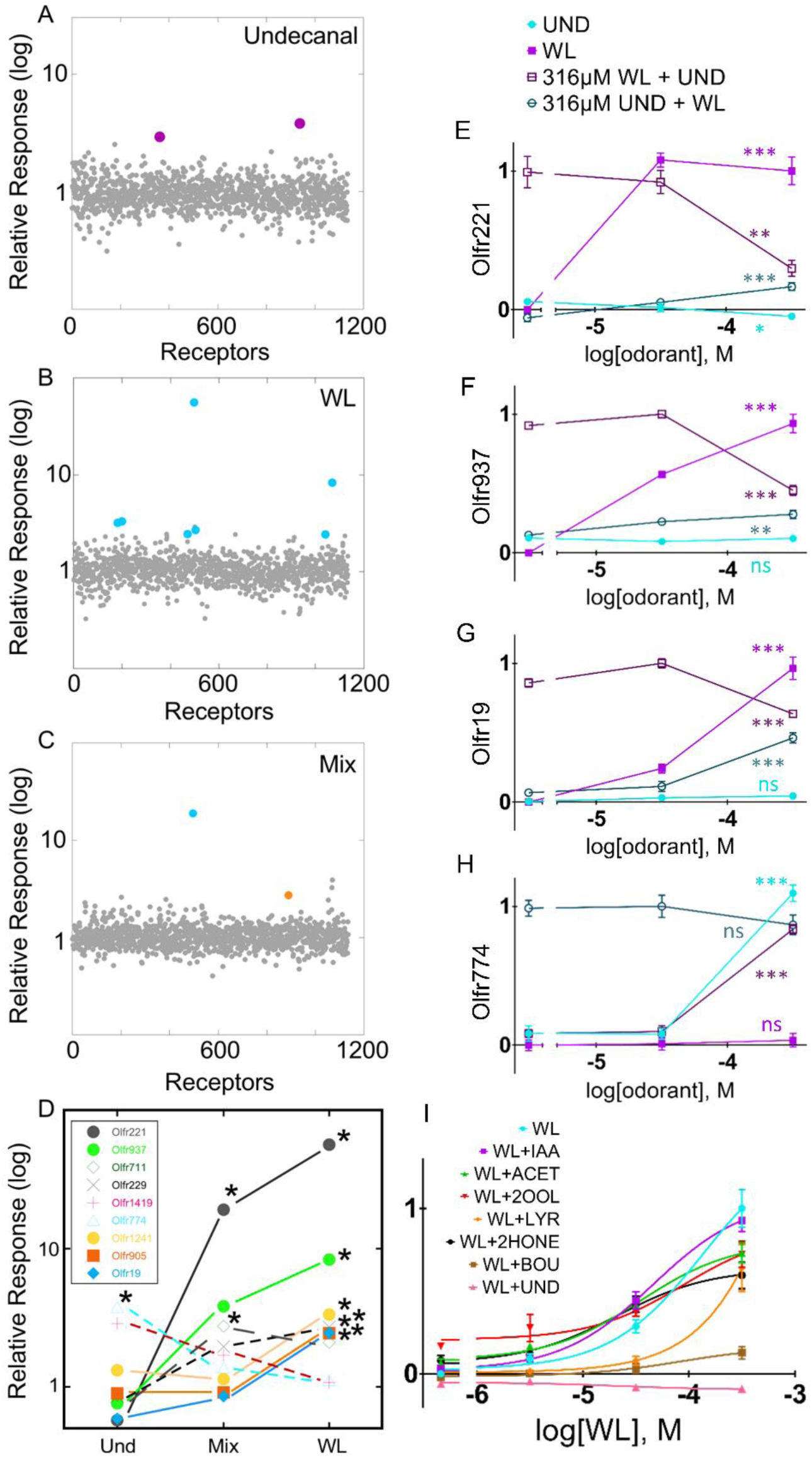
Odorant receptors responsive to undecanal (Und) and whiskey lactone (WL). Of the ORs responsive to either undecanal or whiskey lactone *in vivo* (**A, B**), only Olfr221 and Olfr711, both responsive to whiskey lactone or showing weak evidence of a response to whiskey lactone, respectively, responded to the mixture of undecanal and whiskey lactone *in vivo* (**C**). The magnitude of *in vivo* responses to the mixture were decreased for all but Olfr711 (**D**). *In vitro* measures of responses to undecanal (solid blue), whiskey lactone (solid violet), undecanal mixed with 316µM of whiskey lactone (open violet), and whiskey lactone mixed with 316µM of undecanal (open blue) in panels **E, F, G**, and **H**. Responses of Olfr937 to whiskey lactone alone and to whiskey lactone mixed with seven different odorants at 316 µM (ACET: acetophenone; BOU, bourgeonal; IAA: isoamyl acetate; 2OOL: 2-octanol; LYR: lyral; 2HONE: 2-heptanone) in panel **I**. In the *in vitro* assay plots the x-axes represent the log of the odorant concentration (M) and the y-axes show the normalized Glosensor luminescence response to cAMP. ANOVA trend analysis *p<0.05; **p<0.01; ***p<0.001 in panels **E, F, G**, and **H**.

To determine whether the reduced responses *in vivo* were due to suppressive interactions, we tested several ORs *in vitro*. The strong responses of Olfr221 and Olfr937 to whiskey lactone were both suppressed in a dose-dependent manner by undecanal (Figure 5E, F). Although the undecanal effect on Olfr221 basal activity was a significant dose-dependent trend (Figure 5E), the effect size was so small (15% of basal activity reduction for the highest dose of undecanal) that we do not consider this sufficient evidence to claim that the ability of undecanal to suppress Olfr221 responses to whiskey lactone was due to inverse agonism. Olfr19, which responded less strongly to whiskey lactone, also suffered a dose-dependent decrease in response when increasing concentrations of undecanal were present (Figure 5G). In contrast to the sensitivity of these three whiskey lactone-responsive ORs to undecanal, we found no convincing evidence that Olfr774 was sensitive whiskey lactone. The small decrements in Olfr774 responses to undecanal when high doses of whiskey lactone were mixed with undecanal were not part of a significant dose-dependent trend (Figure 5H).

To further test the hypothesis that interactions between odorants at ORs are common, we selected Olfr937 and tested whether 5 additional odorants would have effects on its responses to whiskey lactone (Figure 5I). As we already demonstrated, undecanal suppressed the ability of Olfr937 to respond to whiskey lactone while isoamyl acetate had no effect on Olfr937. Like undecanal, the structurally related odorants bourgeonal and lyral also suppressed Olfr937 responses to whiskey lactone, with bourgeonal being more effective. The other odorants tested failed to produce a significant change in the dose-response relationship between Olfr937 and whiskey lactone.

## Discussion

With the ability to measure the responses of all mouse ORs and TAARs simultaneously *in vivo*, we identified odorant agonists for 30 ORs: 14 that responded to the headspace air above a solution of 5% isoamyl acetate, 7 to 50% whiskey lactone, 2 to 5% undecanal, and 7 to 2% bourgeonal. We observed no responses from TAARs, consistent with evidence that they are primarily receptors for odorant molecules containing amines (Liberles and Buck, 2006; Johnson et al., 2012; Pacifico et al., 2012). We focused only the certainty achieved when ORs respond in the same way *in vivo* and *in vitro* and leave in doubt those ORs whose responsiveness remains in question due to lack of agreement. Of these 30 ORs identified *in vivo, in vitro* tests confirmed agonist activity at 8 ORs: Olfr183, Olfr213, and Olfr1411 for isoamyl acetate, Olfr221, Olfr937, and Olfr19 for whiskey lactone, Olfr774 for undecanal, and Olfr16 for bourgeonal (Table 1). Agreement between *in vivo* and *in vitro* data minimizes the possibility that *in vivo* responses were due to locally-generated metabolites of the tested odorant (Nagashima and Touhara, 2010; Heydel et al., 2013; Kida et al., 2018), so we consider these 8 agonist-OR relationships to be firmly established. We are also confident about agonist-OR relationships where data from the highly sensitive *in vitro* assay explained novel OR responses to binary mixtures or nearly significant responses *in vivo*. This includes isoamyl acetate responses from Olfr167, undecanal responses from Olfr638, Olfr1019, and Olfr605, and Olfr1420 responses both to undecanal and to bourgeonal (Table 1). The other ORs that reached significance *in vivo* must be considered potential receptors for the tested odorants. In the absence of evidence that these ORs are functional *in vitro*, meaning a positive control response to a different odorant in the *in vitro* assay, we cannot conclude that *in vivo* results represent false positive responses.

**Table 1.**
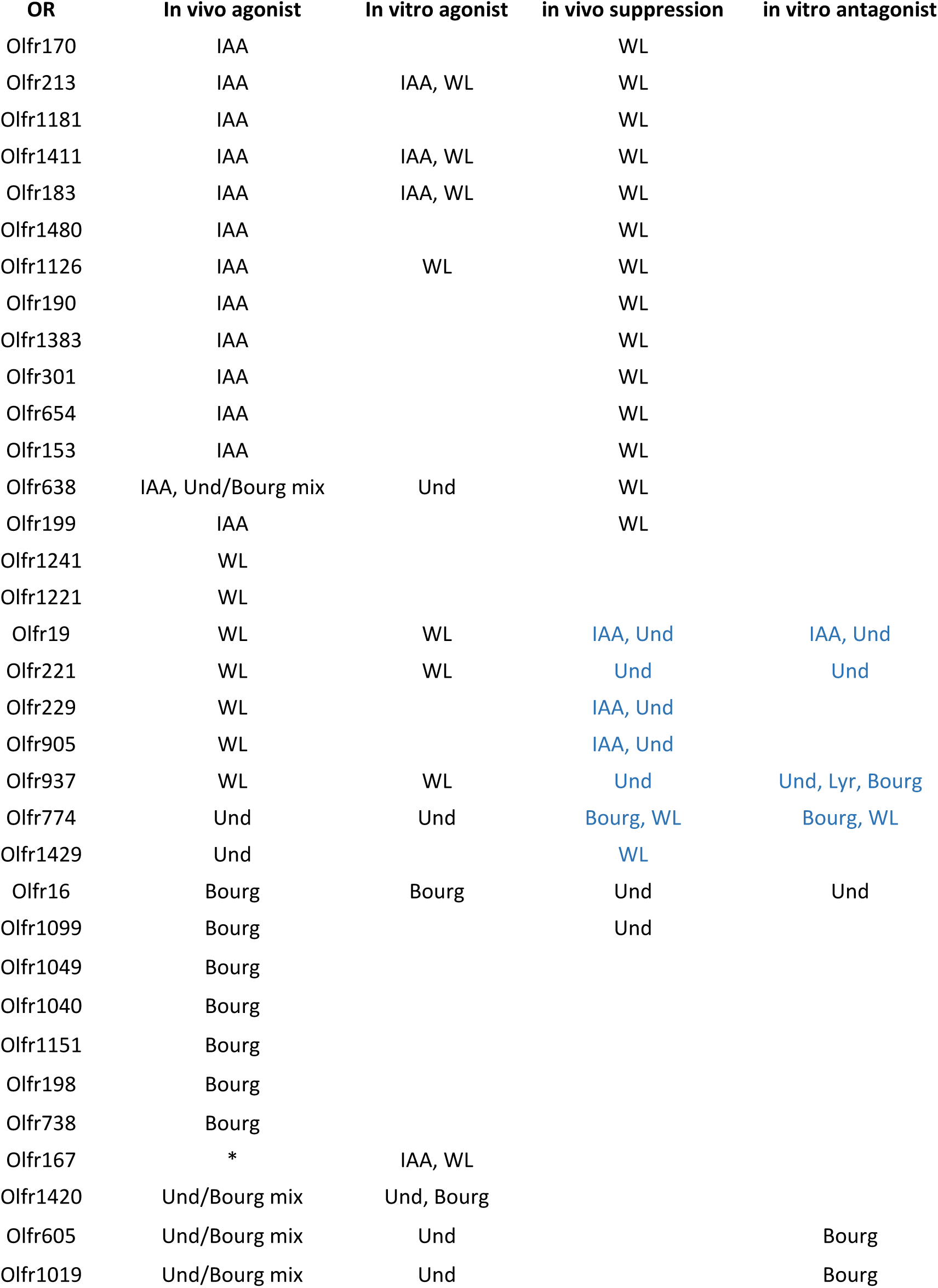
Summary of agonist and antagonist effects detected for ORs that gave significant positive responses to at least one odorant or mixture. Bourg, bourgeonal; IAA, isoamyl acetate; Und, undecanal; WL, whiskey lactone; Lyr, lyral. *, a suspected false negative response to IAA.

| OR | In vivo agonist | In vitro agonist | in vivo suppression | in vitro antagonist |
| --- | --- | --- | --- | --- |
| Olfr170 | IAA |  | WL |  |
| Olfr213 | IAA | IAA, WL | WL |  |
| Olfr1181 | IAA |  | WL |  |
| Olfr1411 | IAA | IAA, WL | WL |  |
| Olfr183 | IAA | IAA, WL | WL |  |
| Olfr1480 | IAA |  | WL |  |
| Olfr1126 | IAA | WL | WL |  |
| Olfr190 | IAA |  | WL |  |
| Olfr1383 | IAA |  | WL |  |
| Olfr301 | IAA |  | WL |  |
| Olfr654 | IAA |  | WL |  |
| Olfr153 | IAA |  | WL |  |
| Olfr638 | IAA, Und/Bourg mix | Und | WL |  |
| Olfr199 | IAA |  | WL |  |
| Olfr1241 | WL |  |  |  |
| Olfr1221 | WL |  |  |  |
| Olfr19 | WL | WL | IAA, Und | IAA, Und |
| Olfr221 | WL | WL | Und | Und |
| Olfr229 | WL |  | IAA, Und |  |
| Olfr905 | WL |  | IAA, Und |  |
| Olfr937 | WL | WL | Und | Und, Lyr, Bourg |
| Olfr774 | Und | Und | Bourg, WL | Bourg, WL |
| Olfr1429 | Und |  | WL |  |
| Olfr16 | Bourg | Bourg | Und | Und |
| Olfr1099 | Bourg |  | Und |  |
| Olfr1049 | Bourg |  |  |  |
| Olfr1040 | Bourg |  |  |  |
| Olfr1151 | Bourg |  |  |  |
| Olfr198 | Bourg |  |  |  |
| Olfr738 | Bourg |  |  |  |
| Olfr167 | * | IAA, WL |  |  |
| Olfr1420 | Und/Bourg mix | Und, Bourg |  |  |
| Olfr605 | Und/Bourg mix | Und |  | Bourg |
| Olfr1019 | Und/Bourg mix | Und |  | Bourg |

Of the ORs responsive to the odorants we tested, only Olfr16 and Olfr1019 had a previously identified agonist. Olfr16 was also shown to be responsive to lyral, an aromatic odorant that is structurally similar to bourgeonal, the strong Olfr16 agonist we discovered (Fukuda et al., 2004). Olfr1019 was also responsive to a fox urine odorant, 2,4,5-trimethylthiazoline, that is structurally dissimilar to the agonist we discovered, undecanal (Saito et al., 2017). For the other responsive ORs in our experiments these are the first odorant agonists identified. The OR response magnitudes we observed suggested that in most cases these odorant agonists were not the best odorant agonists for these ORs, so better agonists for them probably await discovery. Likely exceptions where the identified agonist may be best agonists include isoamyl acetate at Olfr170, whiskey lactone at Olfr221, undecanal at Olfr774, and bourgeonal at Olfr16.

The *in vivo* OR response patterns we detected, often called combinatorial codes (Malnic et al., 1999), usually had a characteristic structure consisting of a few highly responsive ORs and a greater number of weakly responsive ORs. These ORs were rarely related in sequence, but instead came from widely different OR subfamilies (Figure S2). These features were exemplified by the OR response patterns of isoamyl acetate, whiskey lactone, and bourgeonal. We predict that most OR response patterns to an odorant have this structure, a hypothesis supported by the patterns of variation in intensity of glomerular responses to odorants (Friedrich and Korsching, 1997; Wachowiak and Cohen, 2001; Fried et al., 2002; Bozza et al., 2004). Its source must be the forces of natural selection that drive expansion and diversification of OR genes. These evolutionary questions are complicated and will be difficult to address rigorously until we better understand how odor perception is encoded in mammals. We also predict that the strongly responsive ORs in these patterns are key elements most responsible for the perception of an odorant. This hypothesis is supported by odorant-specific decrements in olfactory performance in mice lacking an OR or TAAR strongly responsive to an odorant (Sato-Akuhara et al., 2016; Dewan et al., 2018). It is also consistent with evidence that allelic variants of specific OR genes can result in altered odorant sensitivity (Keller and Vosshall, 2007; Geithe et al., 2017; Dewan et al., 2018).

When encountering odorant mixtures, the ability to perceive individual odorants in the mixture is at risk due to suppression of key ORs by other odorants acting as antagonists, inverse agonists, or partial agonists that compete with full agonists for receptor occupancy. When testing binary mixtures of odorants, we found interactions at ORs to be relatively common even when the odorants were not previously known to interact. *In vivo*, higher delta values are more reliable indicators of response than lower delta values, and ORs not reaching significance cannot be presumed to be unresponsive. For this reason, we claimed that an OR may be non-responsive only if we had other evidence that it does respond in the same assay system. We identified instances where ORs that respond to an odorant presented alone fail to reach significance when this odorant is mixed with another odorant *in vivo*. We tested most of these instances using the *in vitro* assay and claimed confidence that a pharmacological interaction has occurred only when the two approaches agree. Even testing just 3 different binary mixtures involving only 4 odorants, we found evidence of all 3 mechanisms of suppression. We detected a case of an inverse agonist - bourgeonal acting at Olfr577, the second documented case of inverse agonism by an odorant at an OR (Reisert, 2010). More common were the cases of antagonism. Isoamyl acetate was an antagonist at Olfr19. Bourgeonal was an antagonist at Olfr605, Olfr1019, Olfr937, and Olfr774. Undecanal was an antagonist at Olfr16, Olfr221, Olfr937, and Olfr19. We have not yet attempted to determine whether these are competitive or noncompetitive antagonists, but we hypothesize that they will prove to be competitive antagonists. We also demonstrated suppression of a strong odorant agonist by a weak odorant agonist. Bourgeonal was a much weaker agonist for Olfr1420 than undecanal and we observed a dose-dependent decrement in the response of Olfr1420 to 100 µM undecanal as more Olfr1420 proteins became occupied by increasing amounts of bourgeonal. Hypoadditive responses of flies to binary mixtures and the modeling of mammalian OSN response patterns suggest that such partial agonist effects are a common mechanism of interaction between odorants (Munch et al., 2013; Reddy et al., 2018). However, our data indicate that antagonism is the most common form of interaction between odorants at ORs. The prevalence of this variety of suppressive interactions appears to be a common reason why the combinatorial codes of mixtures differ from the simple summation of the OR response patterns of the component odorants. Odorant interactions at ORs cannot help but modulate mixture OR response patterns and could prevent recognition of an odorant’s combinatorial code, even in cases as simple as a binary mixture. These limitations probably help determine whether a mixture is perceived as the odorant elements that comprise the mixture or instead as a unique odor object.

Some of the broadest and strongest suppressive interactions we observed *in vivo* were the effects of whiskey lactone on OR responses to isoamyl acetate. These findings confirm *ex vivo* data from rat electroolfactograms and recordings from isolated rat OSN recordings, which both agree that whiskey lactone suppresses responses to isoamyl acetate (Chaput et al., 2012). We could not confirm this interaction *in vitro* because most ORs specifically responsive to isoamyl acetate *in vivo* instead showed robust responses to both whiskey lactone and isoamyl acetate *in vitro*. These findings exactly parallel *in vitro* heterologous expression data from 2 human ORs, which also responded to both isoamyl acetate and whiskey lactone (Chaput et al., 2012). Our data extend these earlier observations by showing that this *in vivo* versus *in vitro* difference in the action of whiskey lactone extends to the level of individual ORs from the same species. Our *in vitro* results also demonstrate that ORs responsive to isoamyl acetate are not selectively sensitive to low agonist concentrations and therefore fail to respond to the higher concentration resulting from the mixture of two odorant agonists, a phenomenon suggested by observations of glomeruli dropping out of response patterns as concentrations of monomolecular odorants increase (Friedrich and Korsching, 1997; Rubin and Katz, 1999; Meister and Bonhoeffer, 2001; Wachowiak and Cohen, 2001; Fried et al., 2002; Bozza et al., 2004; Jiang et al., 2015). We believe that the *in vitro* data are technically correct in confirming that many ORs responsive to isoamyl acetate are sensitive to whiskey lactone, but that differences in the environments provided by OSNs and cultured cells in some way allow isoamyl acetate-responsive ORs to respond differently to whiskey lactone *in vitro* than they do *in vivo*. The rat electroolfactogram recordings and isolated rat OSN recordings (Chaput et al., 2012) indicate that mixtures of whiskey lactone and isoamyl acetate do not cause abnormal desensitization of ORs or OSN responses, a potential explanation for absence of OR responses *in vivo*. The recordings from isolated OSNs exclude the possible suppressive effects of metabolites of whiskey lactone that might be produced locally in the olfactory mucus or in neighboring cells, as shown for some odorants (Nagashima and Touhara, 2010; Heydel et al., 2013; Kida et al., 2018). The isolated rat OSN recordings (Chaput et al., 2012) also argue that whiskey lactone is not simply acting as a weak agonist *in vivo* to suppress responses to a stronger agonist at these ORs. Furthermore, whiskey lactone is not always an agonist at ORs *in vitro*. For example, it acts as an antagonist, not an agonist, at Olfr774. A mechanistic explanation for how whiskey lactone is able to act differently at ORs responsive to isoamyl acetate when these ORs are expressed in heterologous cells awaits further study.

Our data suggest odorant interactions at ORs might often prove to be similar in humans and rodents. In fact, a reason for choosing to test isoamyl acetate and whiskey lactone was the evidence of similar interaction effects in humans and rats. Both species have difficulty perceiving isoamyl acetate when it is mixed with whiskey lactone (Chaput et al., 2012). In contrast to the strong interactions between isoamyl acetate and whiskey lactone, the ORs most responsive to undecanal and bourgeonal (Olfr774 and Olfr16, respectively), though mildly antagonized by the other odorant, continued to respond strongly to the mixture of these odorants. This predicts that undecanal and bourgeonal should both be readily detectable in mixtures of these two odorants. If human ORs responsive to undecanal and bourgeonal behave similarly to mouse ORs responsive to these odorants, it would be consistent with the ability of humans to perceive both odorants when they are mixed (Brodin et al., 2009). To the degree that such correlations between rodents and humans are common, rodents are a valuable model to investigate the pharmacology of ORs and the effects of odorant interactions on OR response patterns.

In conclusion, we tentatively identified new odorant ligands for 30 ORs, directly confirmed agonists for 8 of these ORs, and provided converging lines of evidence to confirm agonists of 5 additional ORs. We measured *in vivo* OR response patterns that provide insight into the combinatorial codes of OR response patterns the brain uses to perceive and discriminate odors. This is particularly important when odors are mixtures of odorants, which is nearly always the case, because odorant interactions could alter OR response patterns. Indeed, we demonstrated that interactions between odorants at ORs are common. We provided the first demonstration of a partial agonist odorant suppressing responses of an OR to a stronger odorant agonist, the second demonstration of an odorant acting as an inverse agonist at an OR, an example of an additive response of an OR to odorant mixtures, and several examples of odorant antagonism at ORs. In summary, our findings are consistent with the interpretation that the impact of odorant interactions at ORs on odor perception will prove to be substantial.

## Materials and Methods

### Chemicals

Sigma Aldrich (St. Louis, MO) was the source of the odorants isoamyl acetate (#W20553-2), whiskey lactone (W380300), undecanal (#U2202), bourgeonal [3-(4-tert-butylphenyl)propanal] (#CDS001981), and the vehicle used to dilute odorants, dimethyl sulfoxide (#D8418).

### *In vivo* assay

The *in vivo* experiments we performed differed little from published procedures described previously (McClintock et al., 2014). This assay takes advantage of odor-stimulated expression of GFP from the activity-dependent *S100A5* gene locus in the S100a5-tauGFP mouse (The Jackson Laboratory, stock number 6709), a mutant strain in which the coding exons of *S100a5* have been replaced by a sequence encoding a fusion of tau and GFP (McClintock et al., 2014). In this project, the S100a5-tauGFP mice used came from a strain back-crossed for 10 generations against C57BL/6J. All procedures with mice were done according to protocols approved by the Institutional Animal Care and Use Committee of the University of Kentucky. Using fluorescence-activated cell sorting to separately collect GFP+ and GFP-cells from dissociated olfactory epithelia of heterozygous S100a5-tauGFP mice after stimulation with the headspace air flushed from vials containing odor or vehicle solutions, then measuring the amount of every OR mRNA in these samples (Affymetrix Clariom S Arrays) we determined which OR mRNAs were enriched in samples of recently activated OSNs. Because each OSN only expresses a single OR gene, the OR mRNAs enriched in odor-stimulated samples compared to clean air controls must encode the ORs that responded to the odorants tested. We also measure TAAR responses in this assay. We have not published evidence of detection of TAAR responses in this assay but just like OSNs expressing ORs, OSNs expressing TAARs can express GFP in these mice, and just like ORs, each TAAR mRNA has a characteristic distribution between GFP+ OSNs and GFP- OSNs; a basal state that is remarkably stable. For these reasons, we expect this assay to be able to detect responses from TAARs.

To control the odor environment experienced by the mice and allow degradation of GFP evoked by prior odor exposure, each mouse was housed individually in specially designed Plexiglas chambers under a flow of 3.1 l/min of filtered air for 40 hrs. During the last 14 hrs that the mice were in the chambers odors were introduced intermittently for 10 s every 20 min via computer-controlled solenoid valves that diverted the flow of filtered air to flush the headspace from a 50 ml vial containing 5 ml of odorant solution. Control mice simultaneously experienced filtered air flushed through a 50 ml vial containing 5 ml of the odorant diluent (the nonvolatile solvent dimethyl sulfoxide). This intermittent stimulation is designed to minimize the effects of receptor desensitization. The 26 hr half-life of GFP (Corish and Tyler-Smith, 1999) means that once GFP expression is initiated subsequent any subsequent receptor desensitization would fail to prevent capture of activated OSN, further minimizing the consequences of desensitization.

Each sample consisted of 3 mice of one or both sexes, ages 7 – 12 weeks and each set of odor-stimulated mice was paired by litter and sex with a set of vehicle-stimulated mice. At the completion of odor exposure, olfactory mucosae were dissected and cells dissociated in a procedure involving papain, trypsin, deoxyribonuclease, and low calcium saline as described previously (Yu et al., 2005; Sammeta et al., 2007). Cells from 3 identically treated mice were pooled, and fluorescence-activated cell sorting (FACS) was performed in the University of Kentucky Flow Cytometry and Cell Sorting Facility using an iCyt Synergy cell sorting system to collect GFP^+^ and GFP-negative (GFP^-^) cell samples. Total RNA was isolated using the Qiagen RNeasy Micro kit (catalog #74004). RNA quantity was measured using Affymetrix Mouse Clariom S arrays in the University of Kentucky Microarray Facility. The microarray data are available in the Gene Expression Omnibus under the accession numbers GSE123784, GSE123788, GSE123789, GSE123790, GSE123791, and GSE123792. Data were initially processed using Affymetrix GeneChip Command Console software to generate globally normalized quantities for each gene transcript cluster. Additional processing to generate GFP^+^/GFP^-^ ratios from the microarray signal intensities was done in Microsoft Excel. These GFP+/GFP-enrichment ratios help to normalize effect across the different abundances of OR mRNAs and across differences in constitutive activity of ORs. Data for each gene are reported as the relative response (Delta), the GFP^+^/GFP^-^ ratio of signal from odor-stimulated mice divided by the GFP^+^/GFP^-^ ratio of signal from vehicle-stimulated mice. Consider the following four quantities, where a stimulant is an odorant or mixture:

sp: OR mRNA abundance for a stimulant (odor) in GFP+ cells

sn: OR mRNA abundance for a stimulant (odor) in GFP-cells

cp: OR mRNA abundance for the control sample in GFP+ cells

cn: OR mRNA abundance for the control sample in GFP-cells

Delta = (sp/sn)/(cp/cn)

Responsive ORs show a large Delta value because their mRNAs increase in the sp sample while simultaneously decreasing in the sn sample, along with the lack of a similar shift in the cp/cn ratio when the response is specific to odor stimulation.

These experiments were done in a paired design, N = 4, with each sample consisting of 3 mice, and each odor-stimulated group of 3 mice was always paired with a group of 3 mice that were simultaneously exposed to vehicle. The stability of OR GFP^+^/GFP^-^ ratios makes it possible to screen for responsive receptors using a relatively small number of replications. A Bayesian hierarchical model was used to obtain normalized measures of odorant effect, accounting for four sources of variation: (1) basal expression of the receptor, (2) odorant effect, (3) nonspecific/batch effect (correlated changes in both odorant and vehicle control in a paired replicate), and (4) random measurement error. For each odorant effect, the posterior mean divided by the posterior SD provides a measure (Z-statistic) that is approximately normally distributed. Local false discovery rates (FDR) were used to estimate the probability that each receptor is responsive to the odorant under the conservative assumption that the majority of the receptors are not responsive to the odorant (Efron, 2008, 2012). Prior work with positive controls has demonstrated that a false discovery rate of 10% is a suitable level of risk for the identification of responsive receptors (McClintock et al., 2014).

### Cell surface expression

Flow cytometry was conducted to evaluate the cell surface expression of ORs. Hana3A cells were seeded in a 35mm dish (Corning) in Minimum Essential Medium containing 10% fetal bovine serum (FBS) (M10). Lipofectamine2000 (Invitrogen) was used for transfection of plasmid OR DNA. A GFP expression vector (30ng) and RTP1s (300ng) were co-transfected with each OR (1200ng) to monitor transfection efficiency and improve OR trafficking, respectively. About 18-24 hrs post-transfection, cells were incubated in phosphate-buffered saline (PBS) containing 1/400 anti Rho-tag antibody 4D2 (Millipore), 15mM NaN3, and 2% FBS and then washed and incubated with 1/100 phycoerythrin (PE)-conjugated donkey anti-mouse IgG (Jackson Immunologicals). 1/500 7-amino-actinomycin D (7-AAD; Calbiochem), a fluorescent, cell-impermeant DNA binding agent, was added before flow cytometry to identify dead cells and exclude PE labeling of internally located ORs in dead cells from analysis. The intensity of the PE signal among live GFP-positive cells was measured and plotted.

### *In vitro* functional assay

The Glosensor assay (Promega) was used to determine the real-time activity of luciferase in Hana3A cells, as previously described.(Zhang et al., 2017). Briefly, firefly luciferase, driven by a cAMP response element promoter (pGlosensor), was used to determine real-time OR activation levels. For each well of a 96-well plate, 10µg pGlosensor, 5µg RTP1s and 75µg of Rho-tagged receptor plasmid DNA were transfected 18 to 24h before odorant stimulations. Plates were injected with 25µL of Glosensor substrate and incubated 2h in dark, room temperature and odor-free environment. The odorants were diluted at the desired concentration in CD293 media supplemented with copper and glutamine and injected at 25µL into each well. The luminescence is recorded for 20 cycles of monitoring. The activity was normalized to the basal activity and the empty vector responses for each receptor. Final OR response was obtained by summing the responses from all 20 cycles. Dose-dependent responses of ORs were analyzed by fitting a least squares function to the data using GraphPrism. To do statistical analyses of dose-dependent responses of ORs, ANOVA models were fit with orthogonal polynomial contrasts. Tests were then applied to the lowest-order term, representing the presence of a monotonic trend.

## Acknowledgments

We thank Dr. Ken Campbell, Dr. Michael Renfro and Mr. Herb Mefford for technical assistance. This work was supported by grants from NIH R01DC014423 to H.M and R01DC014468 to T.S.M..

## Author contributions

T.S.M. and H.M. designed the project. T.S.M., W.B.T., T.S., and P.B performed in vivo monitoring and statistical analysis. C.A.D.M. performed in vitro monitoring and cell surface expression evaluation. T.S.M., H.M., and C.A.D.M. co-wrote the manuscript. All authors discussed the results and commented on the manuscript.

## Supplementary Materials

**Figure S1.**
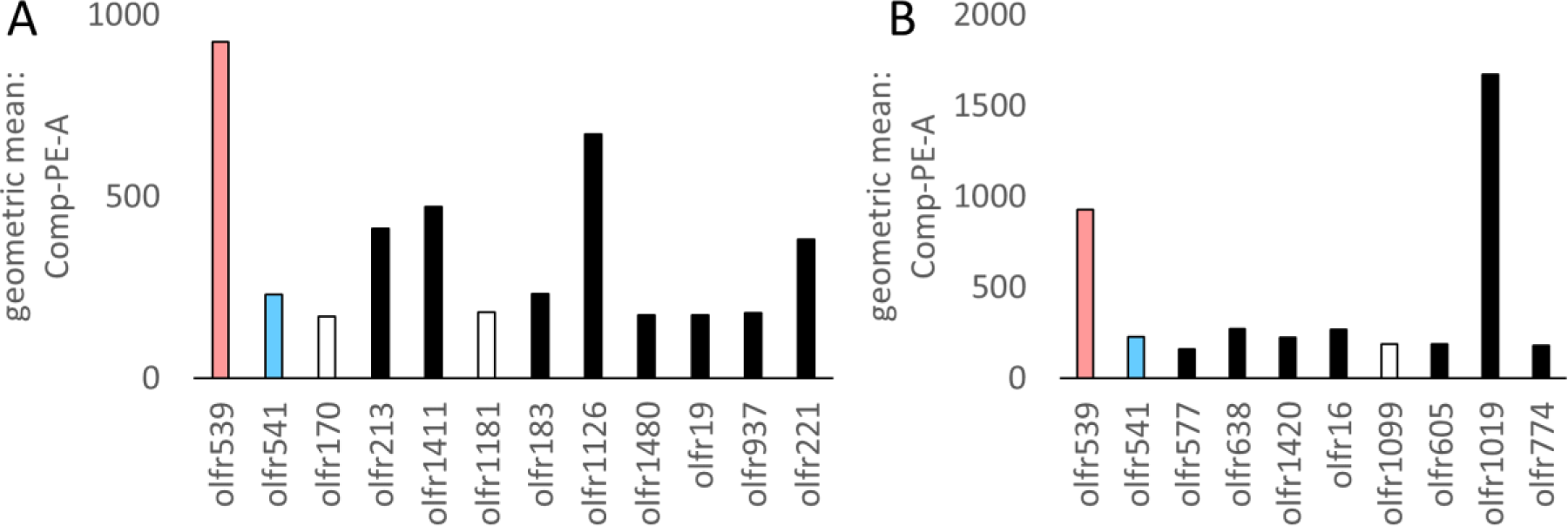
Flow cytometry analysis of levels of receptors in the plasma membrane of live transfected cells. The geometric mean of the Comp-PE-A, witness of cell surface expression, is reported on the y-axis. For both (A) and (B), Olfr539 (pink) and Olfr541 (blue) are positive and negative controls, respectively and ORs responding and not responding *in vitro* are highlighted by black and white filled columns, respectively. (A) ORs responsive to whiskey lactone or isoamyl acetate. (B) ORs responsive to undecanal or bourgeonal.

**Figure S2.**
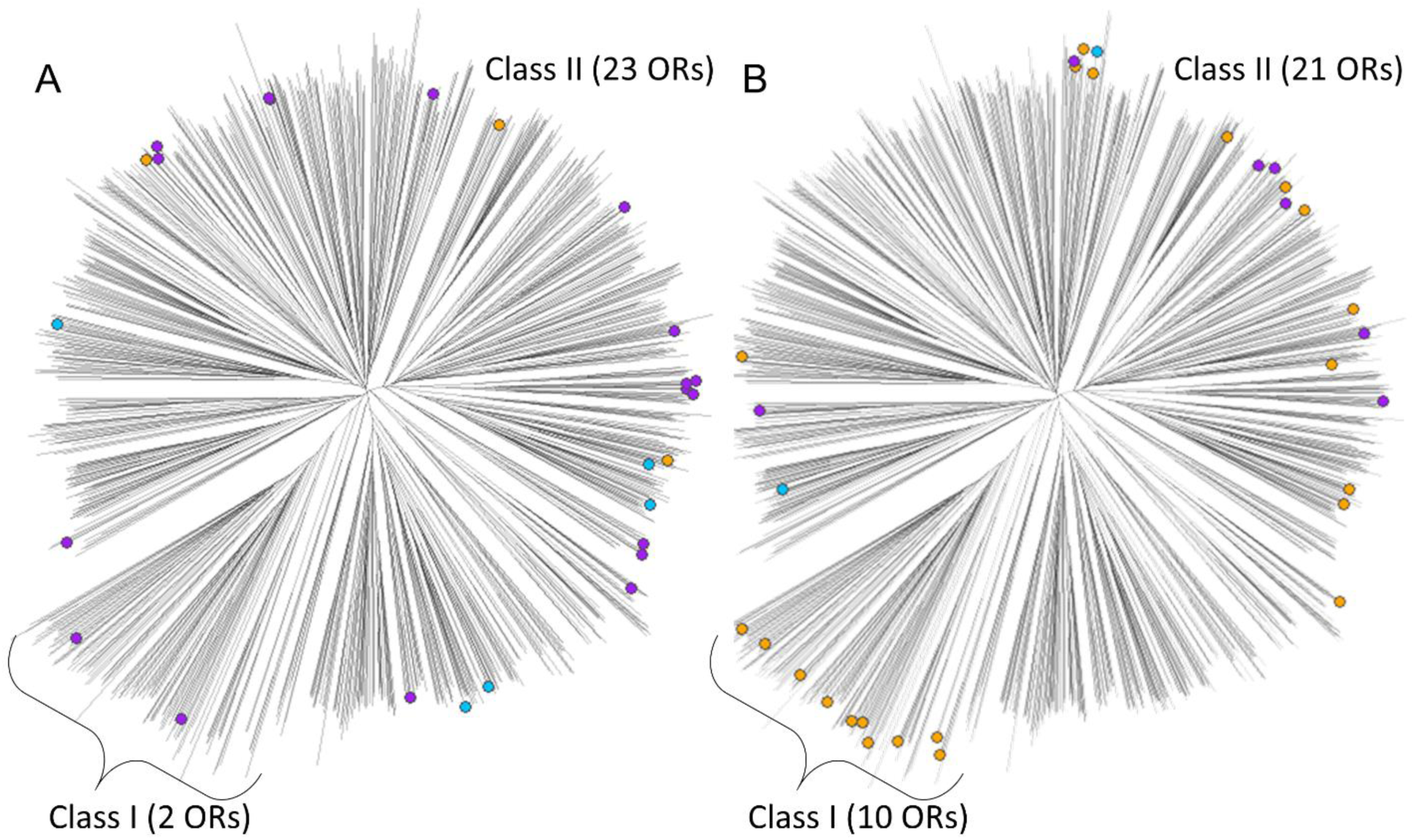
Phylogenetic tree of mouse OR showing both Class I and Class II ORs. ORs responding *in vivo* to whiskey lactone or isoamyl acetate (A) and undecanal or bourgeonal (B) are highlighted by colored dots. (A) ORs responding to whiskey lactone are represented in blue and isoamyl acetate in violet. ORs responding to one of the odorants and still responding to the mixture are represented in orange. (B) ORs responding to undecanal are represented in blue, bourgeonal in violet, and uniquely to the mixture of these two odorants in orange. The number of responding ORs *in vivo* are indicated in parentheses for each OR Class.

**Table S1.** Summary of responses to IAA and WL. Olfr, Entrez gene symbol for each OR tested. mOR, familial name of tested mouse OR. Delta, SD, and FDR results from *in vivo* stimulation by IAA, WL and their mixture. Delta is the ratio of the response to odorant versus response to filtered air as determined by the *in vivo* assay (see methods). SD, standard deviation, N=4. FDR, false discovery rate q-value from the *in vivo* assay. *In vitro* results by Glosensor assay from stimulation done at a single concentration for each stimulus: 1000µM isoamyl acetate (IAA) and 316µM whiskey lactone (WL) and their Mix (1000µM IAA + 316µM WL). nt, not tested. Receptor rows that are represented in bold are those whose *in vitro* dose-response relationships were further determined. *In vivo* and *in vitro* results are colored if the receptor is considered as significantly responding to IAA (violet), WL (blue) or their Mix (orange).

| Olfr | mOR | IAA<br>Delta | IAA<br>SD | IAA<br>FDR | Mix<br>Delta | Mix<br>SD | Mix<br>FDR | WL<br>Delta | WL<br>SD | WL<br>FDR | IAA<br>*in vitro* | Mix<br>*in vitro* | WL<br>*in vitro* |
| --- | --- | --- | --- | --- | --- | --- | --- | --- | --- | --- | --- | --- | --- |
| Olfr170 | mOR273-2 | 9.90 | 0.40 | 0.000 | 4.04 | 0.34 | 0.000 | 1.16 | 0.48 | 1.000 | -1.86 | -0.72 | -1.43 |
| **Olfr213** | **mOR119-3** | **6.14** | **0.40** | **0.000** | **1.82** | **0.34** | **0.659** | **1.22** | **0.31** | **0.941** | **65.35** | **77.68** | **55.32** |
| Olfr1181 | mOR225-9p | 4.06 | 0.35 | 0.000 | 1.81 | 0.47 | 0.910 | 1.72 | 0.37 | 0.652 | 0.71 | 1.42 | 1.79 |
| **Olfr1411** | **mOR208-3** | **3.40** | **0.33** | **0.000** | **2.22** | **0.34** | **0.378** | **1.18** | **0.32** | **0.970** | **49.90** | **45.54** | **46.58** |
| **Olfr183** | **mOR183-2** | **7.13** | **0.56** | **0.000** | **1.31** | **0.75** | **1.000** | **1.21** | **0.36** | **0.957** | **57.48** | **48.26** | **12.52** |
| **Olfr1126** | **mOR264-5** | **2.87** | **0.38** | **0.033** | **1.34** | **0.58** | **1.000** | **0.82** | **0.31** | **1.000** | **8.74** | **11.63** | **9.26** |
| Olfr1480 | mOR202-44 | 3.49 | 0.44 | 0.040 | 2.14 | 0.48 | 0.702 | 1.81 | 0.64 | 0.926 | 1.24 | 16.21 | 13.52 |
| Olfr1383 | mOR256-56 | 2.76 | 0.40 | 0.059 | 1.63 | 0.41 | 0.998 | 1.12 | 0.34 | 1.000 | nt | nt | nt |
| Olfr190 | mOR183-4 | 2.45 | 0.35 | 0.060 | 2.04 | 0.39 | 0.636 | 1.42 | 0.36 | 0.920 | 1.23 | 2.01 | -0.62 |
| Olfr301 | mOR211-8p | 2.67 | 0.40 | 0.068 | 0.97 | 0.55 | 1.000 | 0.78 | 0.41 | 1.000 | 0.66 | -0.72 | -1.99 |
| Olfr654 | mOR38-2 | 2.15 | 0.32 | 0.076 | 1.03 | 0.34 | 1.000 | 1.02 | 0.33 | 1.000 | 0.77 | 1.71 | -1.70 |
| Olfr153 | mOR177-5 | 2.22 | 0.34 | 0.076 | 1.41 | 0.44 | 1.000 | 0.84 | 0.30 | 1.000 | 0.43 | -0.06 | 0.49 |
| Olfr199 | mOR182-14 | 2.34 | 0.38 | 0.088 | 1.98 | 0.45 | 0.756 | 1.18 | 0.34 | 0.970 | 0.56 | 1.14 | 0.74 |
| Olfr638 | mOR5-1 | 2.43 | 0.40 | 0.103 | 1.45 | 0.41 | 1.000 | 1.64 | 0.48 | 0.915 | 1.40 | 2.40 | 0.48 |
| Olfr1384 | mOR256-23 | 2.57 | 0.46 | 0.113 | 0.87 | 0.39 | 1.000 | 1.42 | 0.34 | 0.911 | nt | nt | nt |
| Olfr286 | mOR286-2 | 2.37 | 0.39 | 0.113 | 2.30 | 0.32 | 0.246 | 1.07 | 0.35 | 1.000 | nt | nt | nt |
| Olfr197 | mOR183-3 | 3.16 | 0.52 | 0.117 | 1.22 | 0.50 | 1.000 | 1.03 | 0.38 | 1.000 | 2.60 | 2.97 | -1.21 |
| Olfr1505 | mOR211-4p | 2.06 | 0.34 | 0.128 | 1.11 | 0.40 | 1.000 | 0.67 | 0.49 | 1.000 | nt | nt | nt |
| Olfr167 | mOR272-1 | 2.08 | 0.35 | 0.135 | 1.79 | 0.33 | 0.651 | 1.35 | 0.32 | 0.931 | 56.97 | 51.07 | 20.98 |
| Olfr1241 | mOR231-14 | 1.99 | 0.46 | 0.672 | 0.84 | 0.38 | 1.000 | 3.32 | 0.41 | 0.019 | 0.51 | 1.30 | -0.64 |
| Olfr1221 | mOR233-3 | 2.45 | 0.66 | 0.681 | 0.89 | 0.54 | 1.000 | 3.20 | 0.38 | 0.012 | 0.77 | 1.18 | 0.43 |
| **Olfr19** | **mOR140-1** | **1.31** | **0.39** | **0.891** | **1.41** | **0.36** | **1.000** | **2.45** | **0.35** | **0.091** | **2.26** | **17.21** | **21.07** |
| **Olfr221** | **mOR205-1** | **1.26** | **0.40** | **0.909** | **33.47** | **0.47** | **0.000** | **56.46** | **0.59** | **0.000** | **0.60** | **30.45** | **51.33** |
| **Olfr937** | **mOR171-24** | **0.98** | **0.37** | **1.000** | **4.99** | **0.36** | **0.000** | **8.33** | **0.43** | **0.000** | **-1.47** | **18.51** | **18.15** |
| Olfr905 | mOR167-1 | 1.02 | 0.35 | 1.000 | 1.96 | 0.39 | 0.666 | 2.43 | 0.31 | 0.021 | 0.77 | 3.73 | 2.60 |
| Olfr229 | mOR171-14 | 0.84 | 0.54 | 1.000 | 1.77 | 0.49 | 1.000 | 2.70 | 0.36 | 0.038 | nt | nt | nt |

**Table S2.** Summary of responses to undecanal (UND) and bourgeonal (BOU). Olfr, Entrez gene symbol for each OR tested. mOR, familial name of tested mouse OR. Delta, SD, and FDR results from *in vivo* stimulation by UND, BOU and their mixture. Delta is the ratio of enrichment ratios (see methods). SD, standard deviation N=4. FDR is the false discovery rate q-value from *in vivo* assay. *In vitro* results by Glosensor assay from stimulation done at 316µM UND and 100µM BOU and their Mix (316µM UND + 100µM BOU). nt, not tested. Receptor rows that are represented in bold are those whose *in vitro* dose-response relationships were determined. *In vivo* and *in vitro* results are colored if the receptor is considered as significantly responding to BOU (violet), UND (blue) or their Mix (orange).).

| Olfr | OR | Und<br>Delta | Und<br>SD | Und<br>FDR | Mix<br>Delta | Mix<br>SD | Mix<br>FDR | Bou<br>Delta | Bou<br>SD | Bou<br>FDR | Und<br>*in vitro* | Mix<br>*in vitro* | Bou<br>*in vitro* |
| --- | --- | --- | --- | --- | --- | --- | --- | --- | --- | --- | --- | --- | --- |
| **Olfr774** | **mOR111-11** | **3.76** | **0.45** | **0.029** | **2.96** | **0.33** | **0.001** | **0.73** | **0.35** | **1.000** | **5.03** | **1.58** | **0.39** |
| Olfr657 | mOR40-13 | 0.72 | 0.38 | 1.000 | 2.61 | 0.29 | 0.002 | 0.80 | 0.35 | 1.000 | -0.43 | -0.01 | 0.22 |
| Olfr669 | mOR34-6 | 0.79 | 0.36 | 1.000 | 2.70 | 0.32 | 0.002 | 1.60 | 0.30 | 0.440 | nt | nt | nt |
| **Olfr605** | **mOR202-22p** | **0.63** | **0.31** | **1.000** | **2.91** | **0.36** | **0.003** | **1.01** | **0.34** | **1.000** | **1.35** | **0.67** | **0.36** |
| Olfr1408 | mOR267-4 | 1.45 | 0.31 | 1.000 | 3.08 | 0.37 | 0.004 | 0.80 | 0.30 | 1.000 | -1.65 | -0.27 | 0.41 |
| Olfr594 | mOR32-10 | 0.85 | 0.34 | 1.000 | 2.37 | 0.29 | 0.004 | 0.96 | 0.32 | 1.000 | nt | nt | nt |
| Olfr830 | mOR152-1 | 0.38 | 0.49 | 1.000 | 2.76 | 0.34 | 0.005 | 1.55 | 0.44 | 0.604 | 1.10 | 0.42 | 0.12 |
| Olfr874 | mOR-161-2 | 0.81 | 0.37 | 1.000 | 3.47 | 0.43 | 0.007 | 0.64 | 0.45 | 1.000 | -1.67 | -0.02 | 0.24 |
| **Olfr16** | **mOR23** | **0.68** | **0.31** | **1.000** | **11.03** | **0.82** | **0.008** | **23.44** | **0.61** | **0.000** | **-1.58** | **5.56** | **14.43** |
| **Olfr638** | **mOR5-1** | **2.17** | **0.35** | **0.714** | **3.01** | **0.39** | **0.012** | **0.92** | **0.36** | **1.000** | **1.46** | **0.05** | **0.02** |
| **Olfr1019** | **mOR180-1** | **0.68** | **0.36** | **1.000** | **2.67** | **0.35** | **0.012** | **0.74** | **0.41** | **1.000** | **4.94** | **0.18** | **0.52** |
| Olfr622 | mOR26-1 | 0.88 | 0.49 | 1.000 | 2.18 | 0.29 | 0.025 | 1.00 | 0.45 | 1.000 | -1.38 | 0.24 | -0.10 |
| Olfr884 | mOR162-13 | 0.52 | 0.38 | 1.000 | 2.24 | 0.30 | 0.027 | 0.94 | 0.35 | 1.000 | nt | nt | nt |
| Olfr160 | M72 | 0.79 | 0.31 | 1.000 | 2.18 | 0.31 | 0.032 | 1.39 | 0.36 | 0.669 | -0.71 | 0.57 | 0.75 |
| Olfr1450 | mOR202-33 | 1.22 | 0.46 | 1.000 | 2.20 | 0.33 | 0.042 | 0.81 | 0.36 | 1.000 | -0.12 | 0.07 | 0.43 |
| Olfr521 | mOR101-2 | 1.54 | 0.42 | 1.000 | 2.78 | 0.40 | 0.049 | 1.36 | 0.35 | 0.682 | 0.73 | -0.93 | 0.15 |
| Olfr1002 | mOR175-2 | 1.25 | 0.43 | 1.000 | 2.22 | 0.33 | 0.060 | 0.70 | 0.52 | 1.000 | -0.33 | -1.18 | -0.09 |
| Olfr584 | mOR30-2 | 1.09 | 0.45 | 1.000 | 2.07 | 0.30 | 0.060 | 1.20 | 0.30 | 0.862 | -2.46 | 0.78 | -0.12 |
| Olfr1047 | mOR188-3 | 0.60 | 0.31 | 1.000 | 2.19 | 0.34 | 0.072 | 0.75 | 0.35 | 1.000 | -0.76 | 0.89 | -1.71 |
| **Olfr577** | **mOR7-2** | **0.55** | **0.51** | **1.000** | **1.84** | **0.27** | **0.073** | **1.21** | **0.34** | **0.884** | **-0.04** | **-6.62** | **-5.12** |
| Olfr297 | mOR220-3 | 0.96 | 0.43 | 1.000 | 2.30 | 0.38 | 0.078 | 1.39 | 0.35 | 0.677 | -0.25 | 0.23 | 0.02 |
| Olfr509 | mOR267-14 | 1.00 | 0.32 | 1.000 | 1.92 | 0.28 | 0.080 | 1.16 | 0.30 | 0.915 | -0.42 | 0.94 | -1.86 |
| Olfr510 | mOR204-34 | 1.02 | 0.33 | 1.000 | 1.87 | 0.29 | 0.096 | 1.53 | 0.32 | 0.513 | -1.19 | 0.04 | -2.28 |
| **Olfr1420** | **mOR266-4** | **1.46** | **0.38** | **1.000** | **1.88** | **0.28** | **0.102** | **1.23** | **0.31** | **0.839** | **3.72** | **5.66** | **1.05** |
| Olfr1099 | mOR206-3 | 0.84 | 0.49 | 1.000 | 2.85 | 0.68 | 0.313 | 5.06 | 0.30 | 0.000 | -1.30 | -1.80 | 2.17 |
| Olfr1049 | mOR187-1 | 0.66 | 0.58 | 1.000 | 1.40 | 0.32 | 0.630 | 2.35 | 0.35 | 0.104 | -0.65 | 0.34 | 0.37 |
| Olfr1419 | mOR266-10 | 2.86 | 0.37 | 0.103 | 1.79 | 0.42 | 0.751 | 1.29 | 0.33 | 0.447 | nt | nt | nt |
| Olfr1040 | mOR185-12 | 0.63 | 0.42 | 1.000 | 0.99 | 0.54 | 1.000 | 2.76 | 0.29 | 0.004 | 0.85 | -3.63 | -2.96 |
| Olfr1151 | mOR177-9 | 0.76 | 0.35 | 1.000 | 0.80 | 0.27 | 1.000 | 2.24 | 0.31 | 0.030 | -0.66 | 0.40 | 0.00 |
| Olfr198 | mOR182-8 | 0.83 | 0.43 | 1.000 | 1.03 | 0.28 | 1.000 | 2.21 | 0.30 | 0.032 | -0.52 | 0.56 | -0.16 |
| Olfr738 | mOR106-3 | 0.98 | 0.32 | 1.000 | 0.83 | 0.37 | 1.000 | 2.56 | 0.35 | 0.040 | 0.57 | 0.00 | 0.31 |

**Table S3.**
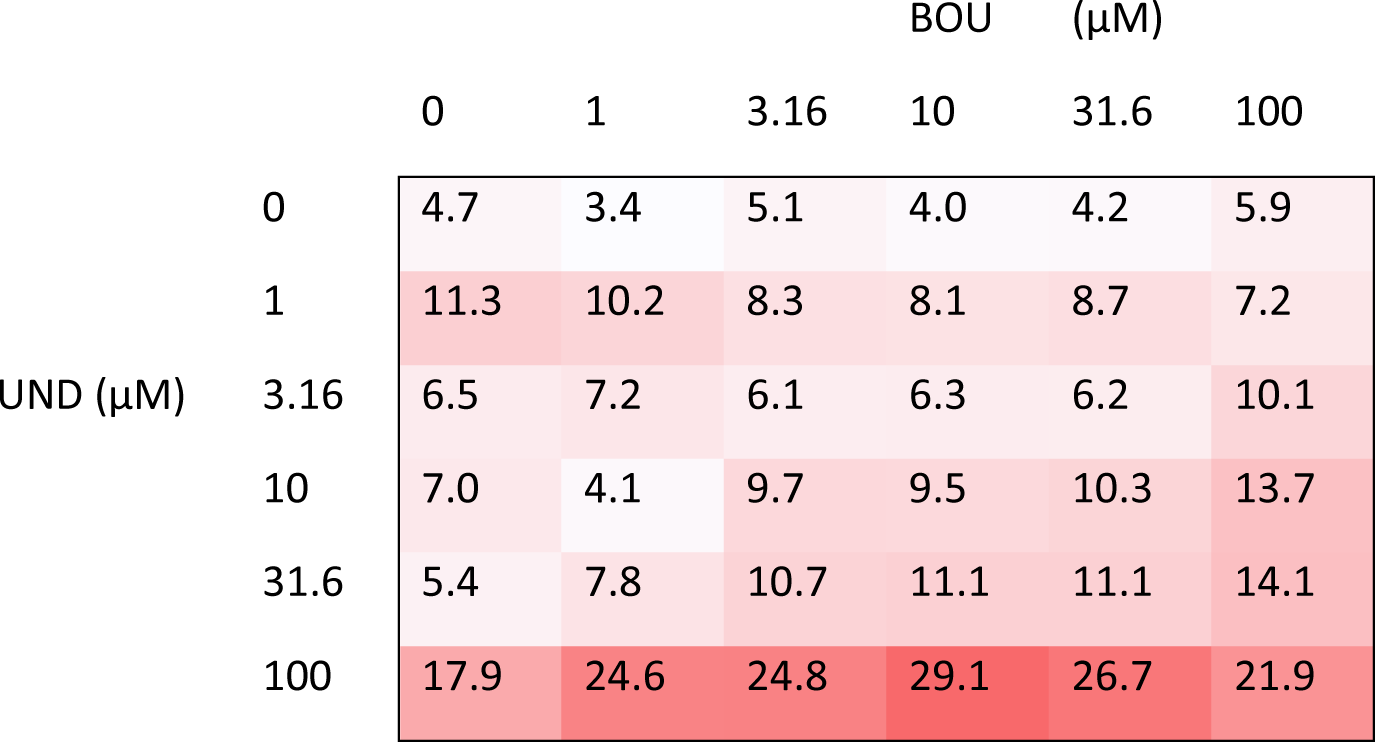
Matrix of Olfr1420 responses to undecanal (UND) and bourgeonal (BOU). Refer to Figure 4D.

**Table S4.**
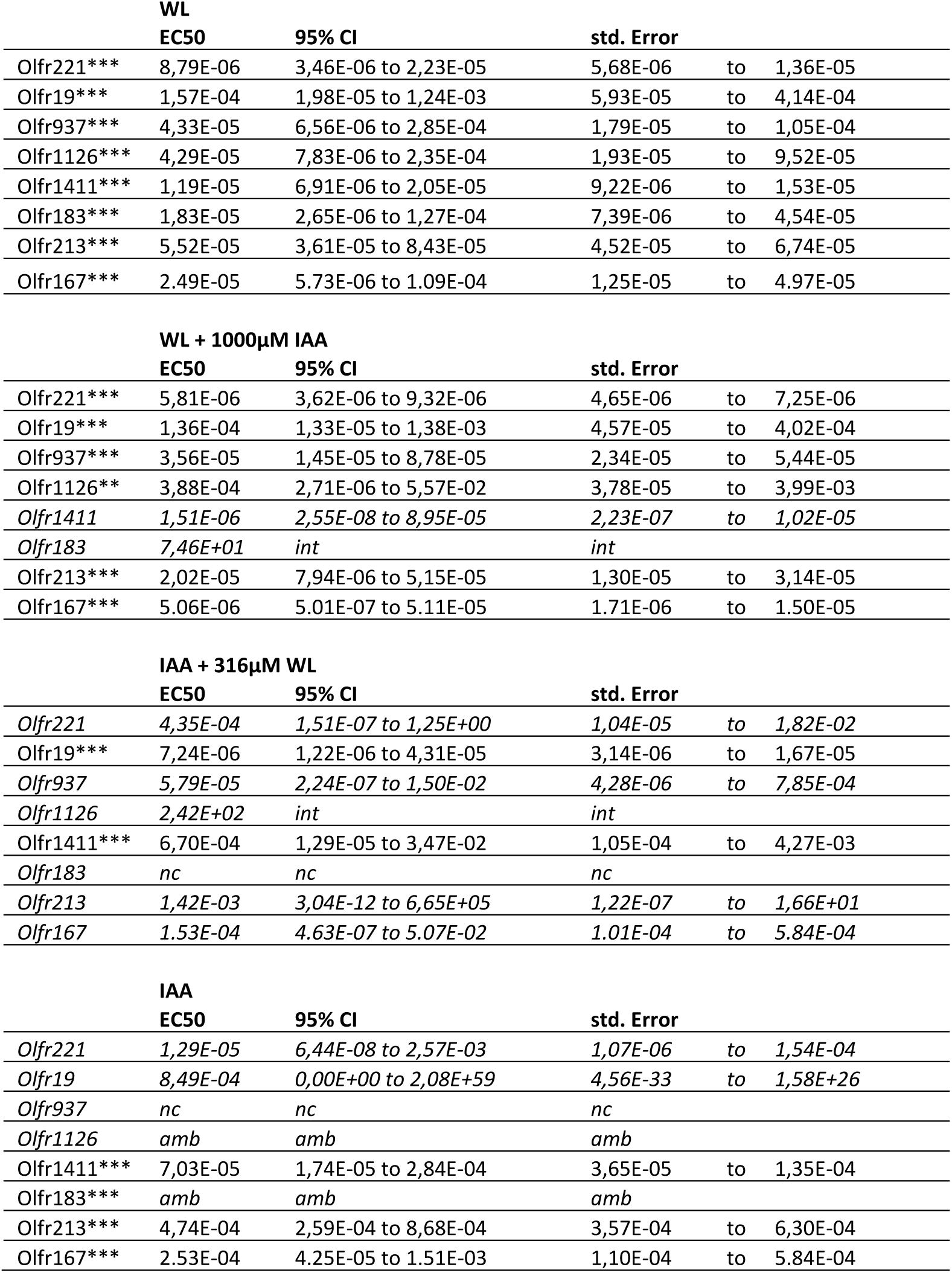
Estimates of EC50 values from dose-response relationship data. These estimates are useful for evaluating the relative efficacy of odorant agonists at ORs and are not meant to be definitive measures of EC50 in each case because not all dose-response relationships achieved saturation. Values are in M for dose-response relationships of whiskey lactone (WL), mixtures (WL + 1000µM IAA and IAA + 316µM WL) and isoamyl acetate (IAA). The 95% confidence interval (95% CI) and the standard error (std. error) are also presented. When dose-response curve fitting is not possible, the value is replaced by *int* for interrupted, *nc* for not converged and *amb* for ambiguous. Non-significant ANOVA trends in dose-response relationships are shown in italics. ANOVA trend analysis *p<0.05; **p<0.01; ***p<0.001.

**Table S5.** Estimates of EC50 values from dose-response relationship data. These estimates are useful for evaluating the relative efficacy of odorant agonists at ORs and are not meant to be definitive measures of EC50 in each case because not all dose-response relationships achieved saturation. Values are in M for dose-response relationships of undecanal (UND), mixtures (UND + 100µM BOU and BOU + 316µM UND) and bourgeonal (BOU). The 95% confidence interval (95% CI) and the standard error (std. error) are also presented. When dose-response curve fitting is not possible, the value is replaced by *int* for interrupted, *nc* for not converged and *amb* for ambiguous. Non-significant ANOVA trends in dose-response relationships are shown in italics. ANOVA trend analysis *p<0.05; **p<0.01; ***p<0.001.

| UND |  |  |  |  |  |
| --- | --- | --- | --- | --- | --- |
|  | EC50 | 95% CI | std. Error |  |  |
| Olfr1019* | 7,98E+01 | *int* | *int* |  |  |
| Olfr1420*** | 6,62E-06 | 3,02E-06 to 1,45E-05 | 4,51E-06 | to | 9,73E-06 |
| Olfr605*** | 1,29E-04 | 6,77E-05 to 2,46E-04 | 9,40E-05 | to | 1,77E-04 |
| Olfr774*** | 4,68E-05 | 1,60E-05 to 1,37E-04 | 2,77E-05 | to | 7,91E-05 |
| *Olfr16* | *2,21E-05* | *2,86E-08 to 1,71E-02* | *8,51E-07* | *to* | *5,73E-04* |
| Olfr638*** | 2,36E-05 | 6,33E-06 to 8,78E-05 | 1,24E-05 | to | 4,48E-05 |
| Olfr577 | *nc* | *nc* | *nc* |  |  |

| UND + 100μM BOU |  |  |  |  |  |
| --- | --- | --- | --- | --- | --- |
|  | EC50 | 95% CI | std. Error |  |  |
| *Olfr1019* | *9,33E-05* | *2,13E-08 to 4,09E-01* | *1,83E-06* | *to* | *4,76E-03* |
| Olfr1420*** | 3,14E-06 | 1,19E-06 to 8,26E-06 | 1,99E-06 | to | 4,95E-06 |
| Olfr605* | *nc* | *nc* | *nc* |  |  |
| Olfr774* | 6,78E-06 | 4,26E+00 to +infinity | 1,38E-06 | to | 3,32E-05 |
| Olfr16* | 6,89E-05 | 1,62E-06 to 2,93E-03 | 1,19E-05 | to | 4,00E-04 |
| *Olfr638* | *4,84E-06* | *0,00E+00 to 4,48E+26* | *4,84E-21* | *to* | *4,84E+09* |
| *Olfr577* | *amb* | *amb* | *amb* |  |  |

| BOU + 316μM UND |  |  |  |  |  |
| --- | --- | --- | --- | --- | --- |
|  | EC50 | 95% CI | std. Error |  |  |
| Olfr1019*** | 3,47E-06 | 6,77E-07 to 1,78E-05 | 1,61E-06 | to | 7,47E-06 |
| Olfr1420* | 2,21E-06 | 9,49E-09 to 5,15E-04 | 1,71E-07 | to | 2,85E-05 |
| Olfr605*** | 1,47E-05 | 4,00E-06 to 5,39E-05 | 7,98E-06 | to | 2,70E-05 |
| Olfr774* | *amb* | *amb* | *amb* |  |  |
| *Olfr16* | *1,17E-06* | *3,47E-09 to 3,96E-04* | *7,64E-08* | *to* | *1,80E-05* |
| *Olfr638* | *amb* | *amb* | *amb* |  |  |
| *Olfr577* | *4,22E+01* | *int* | *int* |  |  |

| BOU |  |  |  |  |  |
| --- | --- | --- | --- | --- | --- |
|  | EC50 | 95% CI | std. Error |  |  |
| *Olfr1019* | *8,30E-05* | *2,20E-13 to 3,13E+04* | *5,81E-09* | *to* | *1,19E+00* |
| Olfr1420*** | 1,00E-04 | 1,26E-05 to 7,94E-04 | 3,66E-05 | to | 2,73E-04 |
| *Olfr605* | *2,56E-06* | *1,26E-07 to 5,22E-05* | *5,96E-07* | *to* | *1,10E-05* |
| *Olfr774* | *nc* | *nc* | *nc* |  |  |
| Olfr16*** | 2,87E-06 | 9,08E-07 to 9,10E-06 | 1,64E-06 | to | 5,02E-06 |
| *Olfr638* | *nc* | *nc* | *nc* |  |  |
| Olfr577* | 5,30E-04 | 8,92E-11 to 3149 | 2,77E-07 | to | 1,01E+00 |

